# Targeting an RNA Editor to Impede H3K27M+ Pediatric Gliomas

**DOI:** 10.64898/2026.05.08.723800

**Authors:** Christian K Ramsoomair, Victoria Alvarez, Felipe Sarmiento, Manuela Aramburu Berckemeyer, Deepa Seetharam, Kajol Kalliecharan, Sreerag N. Moorkkannur, Jonathan Mitchell, Anna Hudson, Justin Taylor, Michele Ceccarelli, Defne Bayik, Oren Becher, Rajeev Prabahakar, Scott M. Welford, Bradley Gampel, Daniel D. De Carvalho, Danny Reinberg, Ashish H. Shah

## Abstract

Diffuse midline glioma (DMG) is a lethal pediatric brain tumor with no curative therapies. Immune checkpoint blockade (ICB) has shown limited efficacy in DMG, largely due to poor T cell infiltration, low immune checkpoint (IC) expression, and a low tumor mutational burden. Here, we identify adenosine deaminase acting on RNA (ADAR), an RNA-editing enzyme that suppresses endogenous dsRNA sensing, as a key mediator of immune evasion in H3K27M-mutant DMG. ADAR is significantly overexpressed in H3K27M tumors relative to wild-type high-grade gliomas, and its depletion selectively suppresses proliferation in patient-derived DMG cells. The H3K27M mutation was found to synergize with ADAR loss to increase retroelement expression, activate type I interferon signaling, and induce immune checkpoint expression. We further identify all-trans retinoic acid (ATRA) as a pharmacologic inducer of ADAR degradation. At clinically relevant doses, ATRA phenocopies ADAR depletion, enhancing antiviral and interferon responses while increasing tumor immunogenicity. In orthotopic immunocompetent DMG models, ATRA enhances CD8+ T cell infiltration and synergizes with ICB and irradiation to improve survival.

## Introduction

Diffuse Midline Glioma (DMG) is the most lethal pediatric brain tumor and despite recent advances in diagnosis and treatment, the majority of children die within 9-12 months of diagnosis with less than 10% surviving 2 years after diagnosis.^1^ Surgical resection is rarely indicated as these tumors predominately occur in essential midline structures of the brain (pons, midbrain, and thalamus). Their diffuse properties and aggressive cancer biology render them resistant to current therapies and standard-of-care radiation therapy is strictly palliative.^2^ Despite recent success in select tumor types, clinical trials with immune checkpoint blockade (ICB) therapy have failed to demonstrate efficacy in high-grade gliomas (HGGs), specifically DMG.^3, 4^ Its scarce lymphocyte infiltration, limited immune checkpoint (IC) expression, and low mutational burden promote an *immunologically cold* tumor microenvironment (TME) that is resistant to ICBs. Thus, there is an urgent need to identify DMG-specific targets that can sensitize the tumor for ICB and other immunotherapies, which has great potential to improve outcomes for this devastating pediatric diagnosis.

A promising strategy termed *Viral Mimicry* has garnered attention for enhancing ICB therapy response rates in *immunologically cold* tumors through the induction of innate antiviral responses.^5–7^ In viral mimicry, dsRNA-sensing immune programs are triggered by aberrant retroelement transcription that produces immunogenic dsRNA species.^8^ Retroelements are genomic relics of ancient viral infections, and their transcription is typically suppressed by heterochromatic epigenetic marks. Viral mimicry therapeutic strategies rely on un-suppressing these retroelement loci, *traditionally* via broad-spectrum epigenetic drugs (inhibitors of DNA methyltransferases or histone deacetylases). This approach boosts dsRNA levels that, in turn, activate downstream type-1 interferon (IFN) responses within the tumor cell, thereby increasing MHC-I and IC ligand expression on the cell surface, and inducing secretion of cytokines that promote CD8+ T-cell activation and ICB therapy sensitization.^8–12^

Herein, we postulated that the driver mutation H3K27M can be leveraged against DMG for viral mimicry activation. The abnormally open chromatin landscape of DMG is one of the most dramatic of all cancers due to its driver mutation: a lysine-to-methionine substitution in the histone H3 tail (H3K27M). This onco-histone effectively acts as an inhibitor of the catalytic subunit, EZH2, of the PRC2 complex, which catalyzes (writes) the heterochromatic H3K27me3 mark. Interestingly, we and others have found that H3K27M results in global decreases of the epigenetic marks known to silence retroelement transcription, including DNA methylation^13, 14^, H3K27me3^13–18^, and H3K9me3^19^. Pharmacologic inhibition of EZH2 has been shown to be sufficient to activate viral mimicry in numerous cancer contexts, including glioblastoma.^20–24^ However, compensatory mechanisms appear to silence these antiviral pathways in DMG. Previous studies have shown that *broad-spectrum epigenetic drugs* activate viral mimicry and significantly improve survival in H3K27M glioma xenografts compared to H3 WT.^13^ However, these drugs have lacked efficacy in DMG clinical trials due to dosing limitations amid systemic toxicities.^25–27^

Despite the dysregulation at the DNA/chromatin-level, viral sensing pathways appear not to be activated due to compensatory mechanisms in DMG. The epi-transcriptome (RNA modifications) of DMG remains largely unexplored, and its potential contribution to the immunosuppressive microenvironment of the tumor is unknown. This gap led us to investigate the therapeutic potential of targeting RNA-level mechanisms.^11^ Such mechanisms promise to be both more efficacious and non-toxic compared to epigenetic drugs since epi-transcriptomic regulators can directly affect the effector molecules of antiviral responses, *i.e.* immunogenic retroelement RNA.^11^

Using an unbiased analysis of epi-transcriptomic regulator expression across pediatric brain tumor proteomic datasets, we identified the adenosine deaminase ADAR as the most significantly upregulated regulator in H3K27M+ gliomas. ADAR is a well-characterized RNA-editing enzyme that destabilizes double-stranded RNA (dsRNA) secondary structures, thereby preventing their recognition by innate immune sensors, such as MDA5, PKR, and OAS2.^28–30^ We demonstrate that ADAR knockdown (KD) in DMG cell lines results in a marked reduction in cellular proliferation, activation of the innate immune response, and increased immune checkpoint expression on the tumor cell surface. Indeed, aberrant retroelement transcription is elevated in DMG as a consequence of K27M-driven epigenetic dysregulation, yet ADAR upregulation mitigates the resulting antiviral response by destabilizing dsRNA intermediates. These findings suggest that DMG cells are dependent on ADAR-mediated RNA editing to evade immune detection. Although there is a dearth of ADAR-specific compounds, we present evidence supporting the use of an FDA-approved compound in a translational approach to enhance the efficacy of immunotherapies *in vivo* against DMG.

## Results

### ADAR is enriched in H3K27M+ pediatric tumors

Given our hypothesis that the epitranscriptome functions as a compensatory mechanism for *viral mimicry* activation in the context of H3K27M-driven epigenetic dysregulation, we leveraged publicly available genetic, transcriptomic, and proteomic datasets from patient-derived cell lines and tumor tissue samples to interrogate this potential relationship.

First, to determine whether an intrinsic signal of suppressed innate immunity is present in H3K27M-positive pediatric gliomas relative to other pediatric brain tumors, we queried the Childhood Cancer Model Atlas (CCMA). Using a previously published interferon-stimulated gene (ISG) signature as a proxy for viral mimicry signal,^31^ we found that ISG scores were significantly elevated in H3K27M-positive pediatric high-grade glioma (pHGG) cell lines compared with those of wild-type pHGGs and other childhood cancers (**Fig. 1a**). Although this finding appears inconsistent with the well-described immunosuppressive features of DMG, analysis of the Pediatric Brain Tumor Atlas (PBTA), which comprises patient biopsy samples containing both tumor cells and components of the tumor microenvironment, revealed no significant differences in ISG scores across pHGG subtypes (**Extended Data Fig. 1a**). Taken together, these results suggest that elevated interferon signaling in H3K27M tumors is primarily tumor cell-intrinsic but constrained by tumor microenvironmental checkpoints that limit baseline viral mimicry activation. We next sought to identify potential epitranscriptomic regulators that could be integral to suppressing this IFN response, such that their functional loss could amplify tumor cell-intrinsic viral mimicry signaling beyond these microenvironmental constraints.

**Figure 1.**
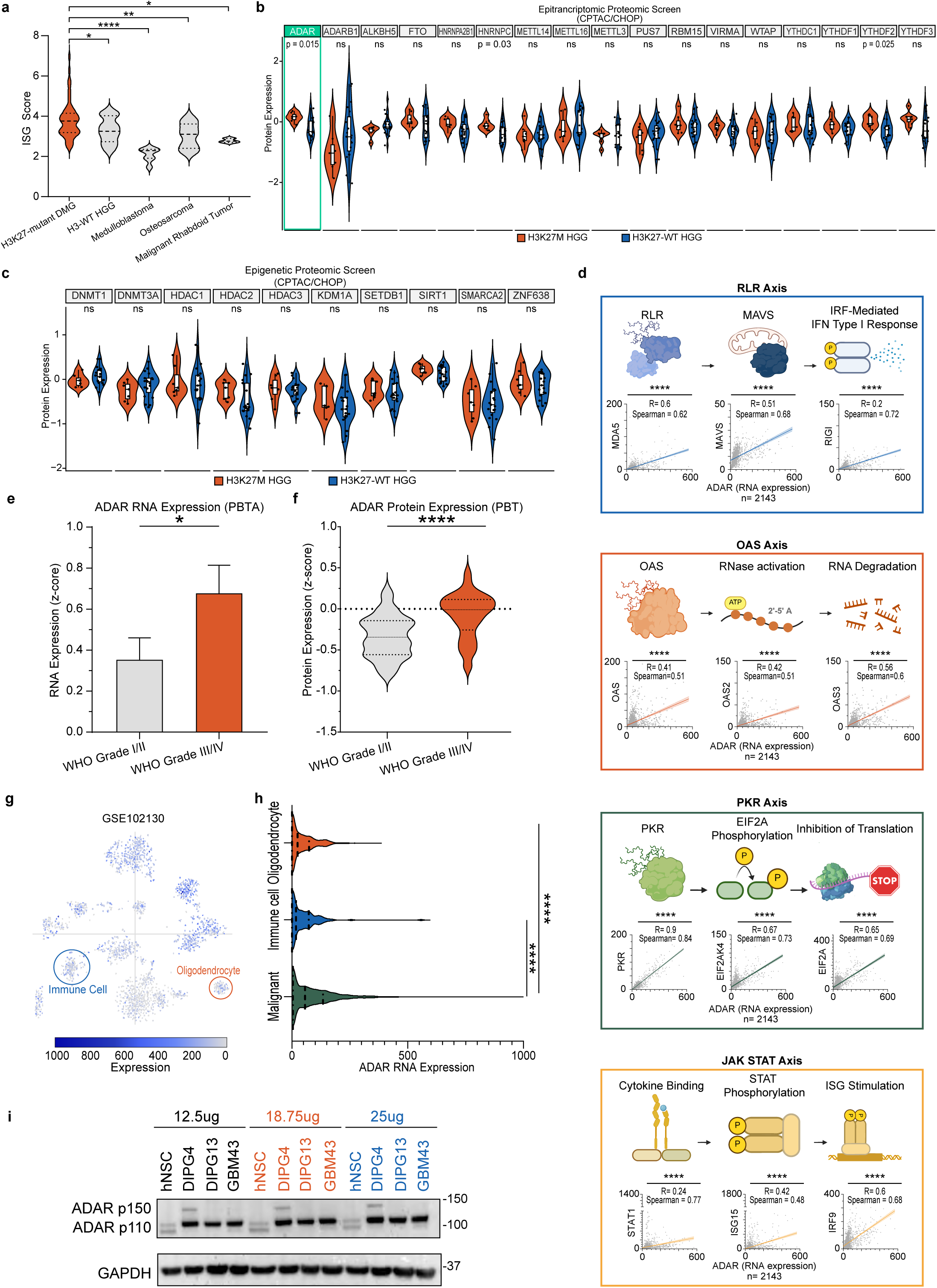
ADAR overexpression is associated with compensatory immune suppression. a) Interferon stimulated gene (ISG) scores in H3K27M-positive high-grade glioma (HGG) cultured cells compared to WT HGGs and other pediatric brain tumors (data from the Childhood Cancer Model Atlas, n=92). b) A proteomic screen comparing expression levels of key epitranscriptomic regulators (n=30) in H3K27M-mutant HGG and WT HGG biospecimens (data from the Clinical Proteomic Tumor Analysis Consortium, n=25). c) A proteomic screen of known immunosuppressive epigenetic regulators (n=10) in H3K27M-mutant HGG and WT HGG biospecimens (data from the Clinical Proteomic Tumor Analysis Consortium, n=25). d) Schematic representations of the RIG-I–like receptor (RLR), JAK–STAT, protein kinase R (PKR), and 2′–5′-oligoadenylate synthetase (OAS) pathways are shown. For each pathway, correlation analyses among three representative core pathway members are shown. e) ADAR RNA expression by tumor grade in pediatric brain tumors (data from PBTA, n=897 pHGG, n=787 pLGG). Error bars represent 95% CI. f) ADAR protein expression by tumor grade in pediatric brain tumors (data from the PBT, n=70 pHGG, n=130 pLGG). g, h) Single-cell RNA sequencing analysis of H3K27M patient samples (GSE102130) delineating differential ADAR expression across malignant cells, oligodendrocytes, and immune cell populations within the tumor microenvironment. i) Immunoblotting analysis of ADAR in DMG/DIPG (SU-DIPGIV, SU-DIPGXIII) and GBM (GBM43) cells, compared with control neural stem cells. (A), (F), and (H) are represented as violin plots where the central dashed line represents the median, upper and lower quartiles are represented by upper and lower dotted lines, respectively. (B) and (C) represented as violin plots with inscribed box and whisker plots where the central line of the box represents the median and the upper and lower quartiles are represented by the top and bottom edges of the box. Data in (E) is presented as means with 95% CI. *P < 0.05, **P < 0.01, ***P < 0.001, ****P < 0.0001 and ns: not significant; Ordinary one-way ANOVA with pairwise t-test FDR corrected for (A), (H). Mann Whitney FDR corrected for (B) and (C). Simple linear regression with 95% CI for (D). Mann Whitney test for (E) and (F).

Using the Clinical Proteomic Tumor Analysis Consortium (CPTAC/CHOP) proteomic dataset, we analyzed patient biopsy mass spectrometry data to exhaustively screen epitranscriptomic regulators and identified adenosine deaminase acting on RNA (ADAR) as the most significantly upregulated factor in H3K27M pediatric high-grade gliomas (pHGGs) compared with wild-type pHGGs (**Fig. 1b**). Verifying this relationship at the genomic level, H3K27M mutant pHGGs also had a significantly elevated percentage of ADAR copy number gain compared to WT (**Extended Data Fig. 1b**). To assess the novelty of this finding, we examined ten well-characterized epigenetic regulators previously implicated in immune suppression and observed no significant differences in expression between mutant and wild-type pHGG samples (**Fig. 1c**).

ADAR, also known as ADAR1, is a well-characterized adenosine-to-inosine (A-to-I) RNA editing enzyme and a member of the three-protein ADAR family.^32, 33^ Two ADAR isoforms are commonly expressed in mammals: p110, which is constitutively expressed and predominantly nuclear, although it can translocate to the cytoplasm upon double-stranded RNA binding; and p150, which is interferon (IFN) inducible and primarily cytoplasmic.^33^ Across multiple cancer types, ADAR has been implicated in regulating innate immune pathways that sense dsRNA and Z-RNA, including RIG-like receptor (RLR), JAK-STAT, PKR, and OAS signaling, whose activation is critical for sensitizing tumors to immunotherapy.^33^ In pediatric brain tumors, we observed strong correlations between ADAR expression and activation of these pathways (**Fig. 1d**), supporting a role for ADAR as a compensatory regulator of innate immunity in diffuse midline glioma. These findings suggest that ADAR upregulation may follow in response to heightened innate immune signaling, functioning to restrain excessive viral mimicry activation. To evaluate ADAR as a potential therapeutic target, we observed strong correlations between ADAR expression and pediatric brain tumor grade across both transcriptomic and proteomic datasets (**Fig. 1e, f**). In addition, single-cell RNA sequencing analysis of H3K27M patient samples (GSE102130) revealed significantly higher ADAR expression in malignant tumor cells compared with normal oligodendrocytes and immune cells within the tumor microenvironment (**Fig. 1g**, **h**). Consistent with these findings, ADAR protein levels were significantly elevated in patient-derived DMG and HGG cell lines relative to nonmalignant, control neural stem cells, as assessed by western blotting (**Fig. 1i**), in agreement with transcriptomic data from the CCMA (**Extended Data Fig. 1c**). We also noted that stem-like DMG cells like SU-DIPG-IV have higher ADAR isoform levels than both DMG cells and astrocytes (hTERT altered) grown in serum-containing media (**Extended Data Fig. 1d**). Collectively, these results support ADAR as a tumor cell–enriched factor associated with glioma aggressiveness and immune suppression in DMG.

### ADAR plays a critical role in the proliferation, migration, and lack of immunogenicity of H3K27M+ cells

After establishing ADAR upregulation in H3K27M+ gliomas and its promising correlative associations with innate immune suppression, we sought to characterize cancer cell fitness upon its knockdown (KD).

Selecting the most efficient candidate from three silencing RNA (siRNA) sequences (**Extended Data Fig. 2a**), we successfully, transiently knocked down ADAR in various DMG and untransformed (hTERT altered) astrocyte cell lines. Utilizing Agilent xCELLigence Real-Time Cell Analysis (RTCA) technology, we observed that in both assayed patient-derived DMG cell lines, SU-DIPG-IV (DIPG4, H3.1K27M) and SF8628 (H3.3K27M), cellular proliferation was markedly reduced (**Fig. 2a**, **b**). Importantly, similarly treated astrocytes showed no phenotypic effects (**Fig. 2c**). To investigate the role of ADAR in self-renewal, we performed an extreme limiting dilution assay (ELDA) using an aggressive patient-derived diffuse midline glioma cell line established at our institution (UM-84), which is capable of robust gliomasphere formation and xenograft generation. At various seeding levels, ADAR KD conditions showed significantly diminished self-renewal capabilities compared to control (**Fig. 2d**, **Extended Data Fig. 2b**). To elucidate the effects of ADAR KD on cell motility and aggressiveness, we performed a transwell migration assay. Again, cells treated with ADAR KD showed a significant reduction in migration compared to control (**Fig. 2e**, **Extended Data Fig. 2c**).

**Figure 2.**
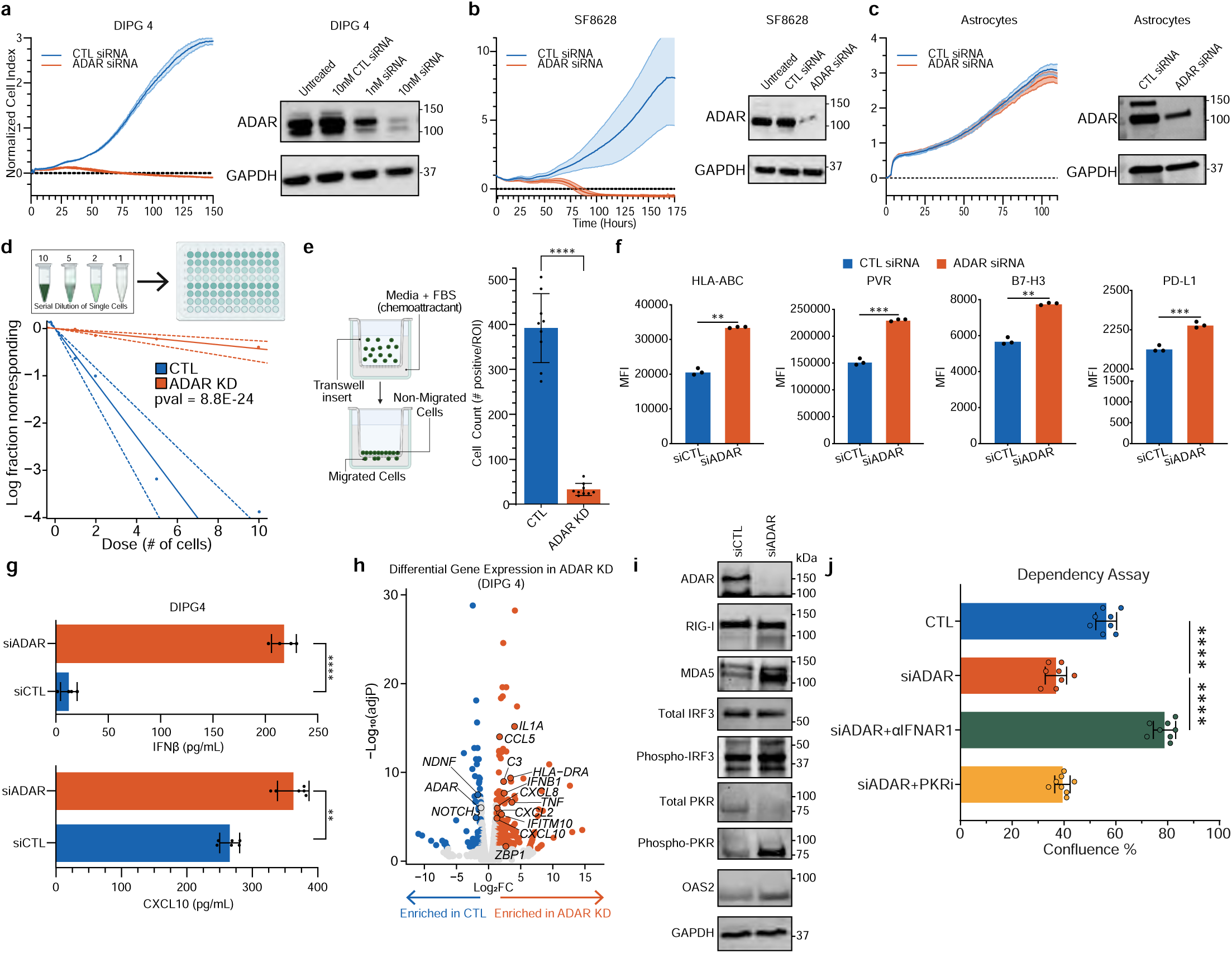
ADAR is critical for H3K27M+ cell proliferation, migration, and immunosuppression. a) xCELLigence proliferation assay with immunoblot confirmation comparing ADAR silencing by RNA interference (siADAR) with negative control treatment (siCTL) in SU-DIPGIV cells. b) xCELLigence proliferation assay with immunoblot confirmation comparing siADAR with siCTL in SF8628 cells. c) xCELLigence proliferation assay comparing siADAR with siCTL in astrocyte (untransformed) cells. d) Extreme limiting dilution assay, with schematic, evaluating gliomasphere-forming capacity and self-renewal of UM-84 cells following siADAR versus siCTL treatment. e) Transwell assay, with schematic, assessing migratory properties of DIPG4 cells following ADAR knockdown compared with control. f) Flow cytometry assessing immunogenic cell surface markers between siADAR- and siCTL-treated SU-DIPGIV cells. g) Enzyme-linked immunosorbent assay (ELISA) measuring concentrations of interferon beta and C-X-C Motif Chemokine Ligand 10 (CXCL10) in siADAR-versus siCTL-treated SU-DIPGIV cells. h) Volcano plot illustrating key differentially expressed genes in DIPG4 following ADAR knockdown. i) Immunoblot analysis of PKR, RIG-I–like receptor, and OAS pathway activation following siADAR or siCTL treatment. j) Dependency rescue assay following ADAR knockdown using type I interferon receptor blockade and PKR inhibition to identify pathways required for the anti-tumor response in SU-DIPGIV cells. (E),(F),(G),(J) is presented as mean ± SD;*P < 0.05, **P < 0.01, ***P < 0.001, ****P < 0.0001 and ns: not significant; Chi-Squared test for (D); Mann Whitney test for (E) and (G), Welch’s two-tailed unpaired t-test for (F), and ordinary one-way ANOVA with pairwise Mann Whitney FDR corrected for (J).

To characterize the role of ADAR in immune suppression of H3K27M+ cells, we first sought to assess immunogenicity on the tumor cell surface via flow cytometry. ADAR KD resulted in consistent and significant upregulation of antigen presentation machinery and IC ligands compared with siCTL, including HLA-A/B/C, PVR (CD155), B7-H3, and PD-L1 (**Fig. 2f**). We next asked whether this increase in immunogenicity induces signaling pathways and secreted factors known to promote activation of the adaptive immune system, particularly T cells. Of translational importance, ADAR KD cells showed significantly elevated concentrations of type 1 interferon (IFNβ) and T-cell activating chemokines (CXCL10) in the culture supernatant via enzyme-linked immunosorbent assay (ELISA, **Fig. 2g**).

Consistent with evidence from other cancers and our correlation analyses in **Fig. 1f**, RNA sequencing revealed the top differentially expressed genes upon ADAR KD to be predominantly inflammatory interferon- or chemokine-related genes (**Fig. 2h**). Furthermore, immunoblot analysis showcased significant activation of dsRNA sensing such as protein kinase R (PKR), RLR, and OAS pathways (**Fig. 2i**). To assess whether the phenotypic effects of ADAR KD are dependent on these pathways, we blocked IFN-α/β receptor 1 (IFNAR1) using antibodies and inhibited PKR pharmacologically, thereby preventing aberrant activation of RLR and PKR signaling, respectively. MDA5 and other RLRs are known drivers of type I interferon production, while PKR activation inhibits protein synthesis with growth arrest. Interestingly, IFNAR1 blockade significantly rescued viability upon ADAR KD while PKR inhibition could not (**Fig. 2j**, **Extended Data Fig. 2d**). Thus, RLR-mediated IFN signaling appears to be more important in its anti-tumor response of ADAR KD than PKR. For context, previous studies have demonstrated that ADAR KD in oral squamous cell carcinoma is rescued by depletion of any of the following: IFNAR1, STING, MAVS, or PKR depletion.^31^ To assess long-term adaptation to ADAR depletion, we generated CRISPR knockout (KO) DIPG4 cell lines and observed differential expression of several key viral mimicry regulators. MDA5, OAS2, and MAVS appeared reduced while RIG-I and PD-L1 expression were elevated (**Extended Data Fig. 2e**). Of note, the ADAR isoform p110 appeared to be markedly reduced relative to isoform p150 after CRISPR KO. Collectively, these findings suggest that ADAR is not only a key immunosuppressive regulator in DMG, but also an important driver of tumor aggressiveness and motility.

### ADAR-mediated immune suppression of H3K27M+ cells is highly associated with retroelement repression

Pathway enrichment analyses of RNA sequencing data revealed immune response pathways were the most significantly upregulated following ADAR KD, consistent with activation of viral mimicry programs. Gene Ontology (GO) analysis revealed significant enrichment of inflammatory and defense response pathways (**Fig. 3a**), while Kyoto Encyclopedia of Genes and Genomes (KEGG) analysis highlighted activation of TNF and NF-kappa B signaling pathways (**Fig. 3b**). Additionally, Reactome analysis showed increased signaling by interleukins and decreased pathways associated with cellular migration like extracellular matrix organization (**Extended Data Fig. 3a**). Moreover, key differentially expressed genes included cytoskeletal regulation and immune signaling (**Fig. 3c**). Together, these findings raised the possibility that loss of ADAR unmasks endogenous nucleic acid signals, prompting us to examine the role of retroelement de-repression in driving this immune activation.

**Figure 3.**
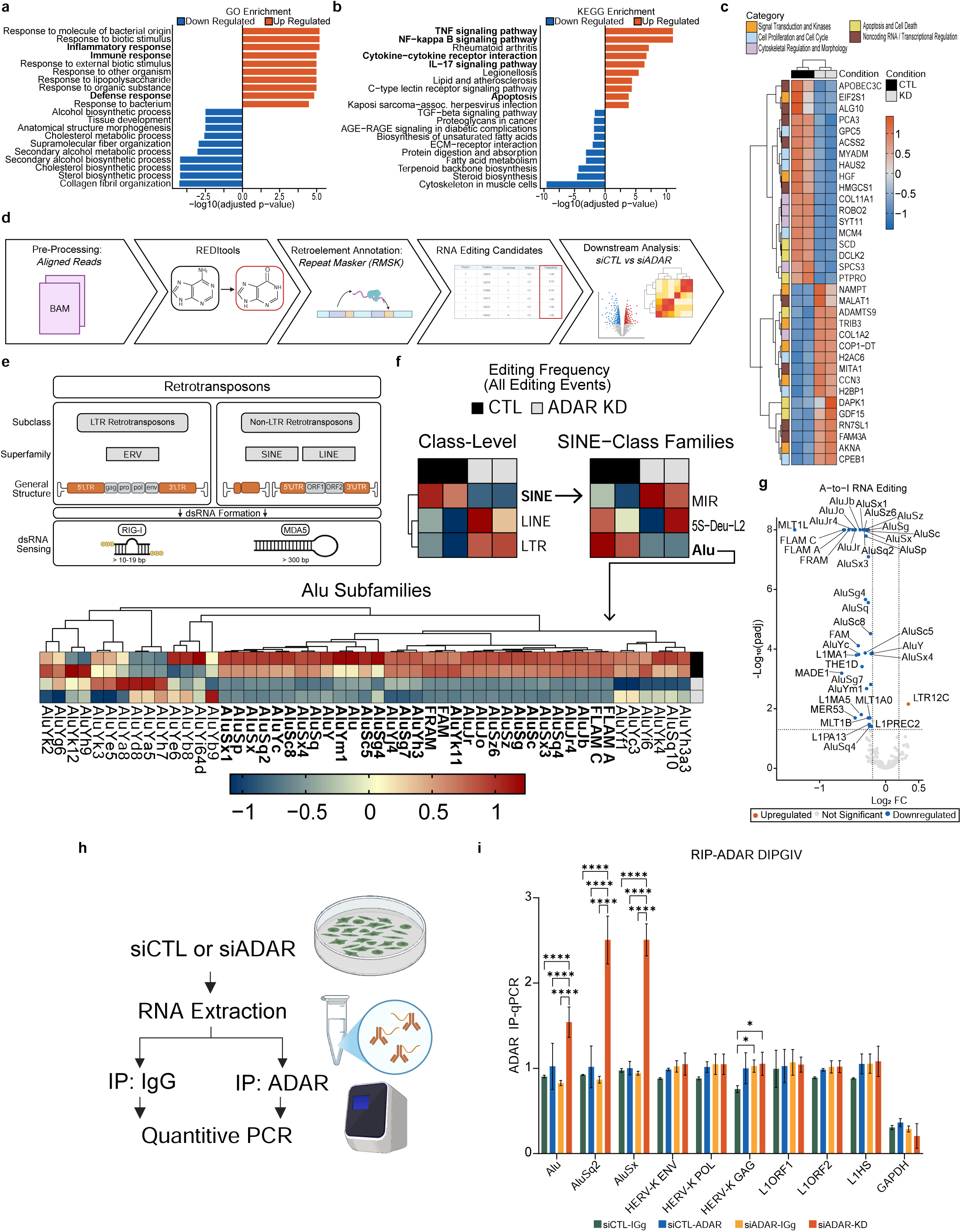
A-to-I editing of Alu/SINE-associated dsRNAs associated with immune suppression in DMG. a) Most significantly upregulated and downregulated Gene Ontology (GO) terms after siADAR treatment relative to siCTL. b) Most significantly upregulated and downregulated Kyoto Encyclopedia of Genes and Genomes (KEGG) terms after siADAR treatment relative to siCTL. c) Heatmap showing selected top differentially expressed genes upon siADAR treatment compared to control. d) Schema illustrating bioinformatic pipeline quantifying A-to-I editing individual retroelement at locus-level between siADAR and siCTL. e) Schematic of retrotransposons by class and family. f) Heatmap shows differentially A-to-I edited retroelement transcripts at Class, Family, and Sub-family levels. g) Volcano plot of differentially A-to-I edited retroelement transcripts ADAR KD compared to control. h) Graphical schematic of RNA immunoprecipitation followed by qPCR (RIP–qPCR). i) RIP–qPCR using anti-ADAR antibodies to quantify enrichment of multiple retroelement species after ADAR knockdown relative to control. (I) is presented as mean ± SD;*P < 0.05, **P < 0.01, ***P < 0.001, ****P < 0.0001 and ns: not significant; Mann Whitney test for (I).

Previous reports have demonstrated that ADAR primarily regulates immunosuppression through the editing and destabilization of Alu/SINE-associated dsRNAs.^28–30^ Leveraging a custom bioinformatic pipeline utilizing REDItools combined with retrotransposon annotation, we quantified levels of differential total and A-to-I RNA editing between ADAR KD and control samples (**Fig. 3d**).^34^ Broadly probing all classes and families of retrotransposons (**Fig. 3e**), we found significantly reduced A-to-I editing within the SINE class, specifically in Alu repeat elements (**Fig. 3f**). Results were verified by qPCR (**Extended Data Fig. 3b**). Importantly, bioinformatic analysis also revealed that the most significantly ADAR-edited Alu/SINEs in DMG are from the oldest- or intermediate-age Alu lineages (FLAM, AluJ, and AluS subfamilies), which have no evidence of pathologic retrotransposition activity (**Fig. 3g**).^35^ To confirm that Alu elements are directly interacting with ADAR, we performed RNA immunoprecipitation (RIP) using anti-ADAR antibodies combined with qPCR. Indeed, we observed enriched Alu retroelement signal under ADAR KD conditions, suggesting both direct binding to ADAR and upregulation upon ADAR depletion (**Fig. 3h**, **i**). Together, our results indicate that ADAR functions as a key immunosuppressive regulator in DMG through its repression of Alu retroelement levels.

### Synthetic immunogenicity induced by combined H3K27M-mediated epigenetic dysregulation and ADAR Knockdown

Given the extensive mutational heterogeneity of DMG, we sought to directly demonstrate that the H3K27M mutation confers a specific sensitivity to ADAR-mediated viral mimicry. We utilized an inducible expression system in 293 T-REx cells engineered in our laboratory^16^ (**Fig. 4a**), which recapitulates the global loss of H3K27me2/3 observed in H3K27M-positive gliomas (**Fig. 4b**). Seventy-two hours after doxycycline induction with or without siADAR treatment, cells were harvested and subjected to downstream analysis. RNA sequencing and bioinformatic analyses revealed distinct transcriptomic states between conditions, yielding several key observations (**Fig. 4c**). First, the top pathways upregulated following ADAR KD in the H3K27M context were related to innate immune signaling and viral mimicry (**Fig. 4d**, **e**). In contrast, these pathways were insignificantly activated after ADAR KD in the uninduced wild-type context. Consistent with this pattern, the majority of the top upregulated genes in ADAR KD H3K27M-induced cells are established regulators of *viral mimicry* and inflammatory interferon signaling (**Fig. 4f**, **g**). Again, we observe that this upregulation of innate immune regulators is absent in the ADAR KD H3 WT condition. Selective activation of OAS and IFN-related pathways in the ADAR KD H3K27M setting were further validated by western blotting and qPCR (**Fig. 4h**, **Extended Data Fig. 4a**). Notably, although MDA5 was among the top 25 upregulated transcripts following ADAR KD H3K27M, its protein levels remained relatively unchanged by western blot. Similarly, RIG-I was among the top 10 upregulated mRNAs upon ADAR KD H3K27M; however, protein-level analysis revealed a more complex regulatory pattern. Western blot demonstrated suppression of RIG-I following siADAR in both wild-type and K27M contexts, although overall RIG-I levels were higher in K27M in general. In summary, the combination of H3K27M-driven epigenetic dysregulation and ADAR loss induces post-translational regulation of a subset of viral mimicry regulators. Notably, this effect is specific to H3K27M-mutant cells and is not observed in wild-type cells. Consistent with this, ELISA showed significantly increased secretion of type I interferon and the T cell–activating chemokine CXCL10 exclusively in ADAR knockdown H3K27M cells (**Fig. 4i**), further supporting activation of the RLR–MAVS axis and a robust viral mimicry–driven innate immune response in the K27M context.

**Figure 4.**
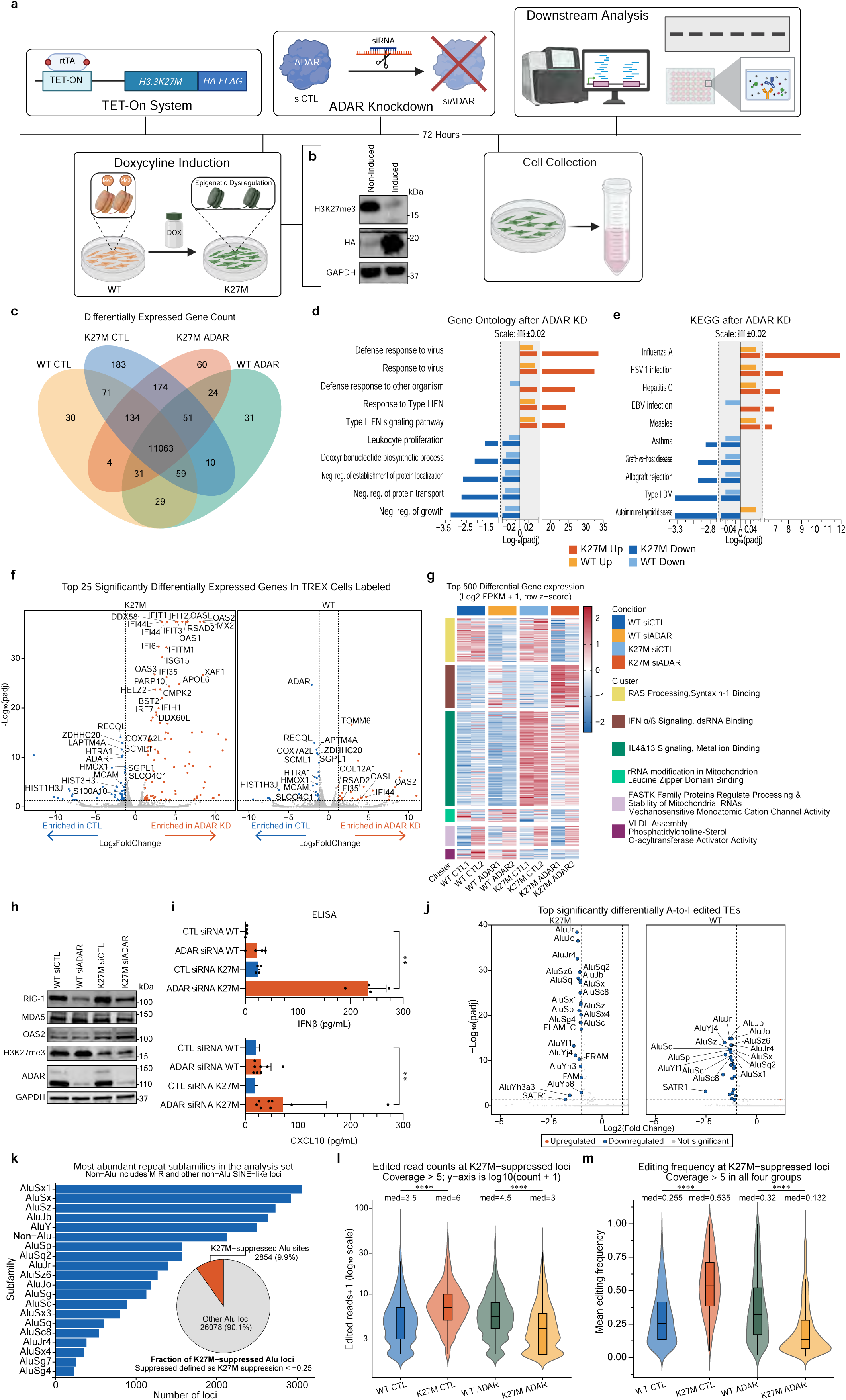
ADAR knockdown enables synthetic immunogenicity in an inducible H3K27M+ model. a) Experimental schema of inducible H3K27M expression platform with ADAR knockdown to evaluate mutation-dependent immunogenic responses. b) Immunoblot validation of epigenetic dysregulation in inducible H3K27M expression platform. c) Venn diagram illustrating differentially expressed genes between conditions. d) GO pathway analysis in H3K27M versus WT context comparing ADAR knockdown to control. e) KEGG pathway analysis in H3K27M versus WT context comparing ADAR knockdown to control. f) Faceted volcano plot comparing significantly differentially expressed genes upon ADAR knockdown in both K27M and WT context. g) Heatmap of the top 500 differentially expressed genes. Samples are hierarchically clustered by condition, and genes are grouped into six distinct expression clusters based on shared expression patterns, with cluster annotations derived from Enrichr. h) Immunoblot-based validation of pathway activation through analysis of representative downstream effectors. i) Type I interferon and CXCL10 levels in culture supernatants measured by ELISA across the four experimental conditions. j) Faceted volcano plot comparing significantly differentially expressed retroelements upon ADAR knockdown in both K27M and WT context. k) Bar plot and nested pie chart illustrating the subfamily abundance and fraction of Alu loci that have enriched ADAR-editing under K27M conditions (K27M-suppressed Alu loci). l, m) Edited read counts and editing frequency of K27M-suppressed Alu loci for all conditions. (I) is presented as mean ± SD; *P < 0.05, **P < 0.01, ***P < 0.001, ****P < 0.0001 and ns: not significant; ordinary one-way ANOVA for (I); Mann Whitney test for (L) and (M). (L) and (M) represented as violin plots with inscribed box and whisker plots where the central line of the box represents the median and upper and lower quartiles are represented by the top and bottom edges of the box.

Using the same custom RNA editing bioinformatic pipeline as above, we quantified differential A-to-I editing between conditions. Remarkably, we observed a striking difference in significant A-to-I editing events between K27M and WT contexts upon ADAR loss (**Fig. 4j**). Again, the most significant changes in editing events upon ADAR loss occurred in Alu repeat elements. By qPCR, we confirmed that total levels of Alu were also significantly upregulated in K27M-induced cells treated with siADAR compared to other conditions. To better understand the basis of this divergence, we next performed a locus-level analysis of Alu editing dynamics across conditions. For each Alu site, we quantified the ADAR-dependent change in editing (*siADAR* − *siCTL*) and compared this response between WT and K27M contexts. We defined *K27M suppression* as the difference in ADAR KD response between epigenetic states:

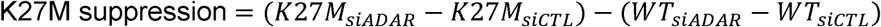

This analysis revealed a subset of Alu loci in which ADAR-mediated editing was markedly elevated at baseline (siCTL) K27M relative to WT (**Fig. 4k**). Notably, nearly 10% of all Alu loci exhibited at least a 25% greater difference in editing between siADAR and siCTL conditions in the K27M context compared with WT. We defined these as K27M-suppressed sites, corresponding to loci where a significant increase in editing occurred following K27M induction and was markedly attenuated upon ADAR KD. Visualization of edited read counts and editing frequencies across conditions suggests that these loci are more actively transcribed within the globally euchromatic landscape associated with K27M (**Fig. 4l**, **m**). In this context, ADAR-mediated editing may function as a compensatory mechanism to constrain dsRNA levels and limit activation of viral mimicry pathways. Collectively, these data suggest that K27M-associated epigenetic dysregulation induces a state of synthetic immunogenicity upon ADAR depletion.

### High-dose ATRA effectively degrades ADAR to phenocopy ADAR KD effects in DMG cells

Despite evidence describing ADAR as a potent immune activator across multiple cancer contexts,^36–40^ no FDA-approved ADAR inhibitors currently exist. Moreover, we were unable to identify any ADAR-targeting compounds in development with demonstrated blood-brain barrier penetration. Recently, however, Li et al. reported the repurposing of all-trans retinoic acid (ATRA) as an ADAR degrader via the ubiquitin–proteasome–mediated pathways.^38^ Importantly, they demonstrated that ATRA treatment increased PD-L1 expression in pancreatic and breast cancer models and synergized with ICIs *in vivo*, as well as in a pilot clinical trial (NCT05482451). Based on these findings, we sought to determine if ATRA may effectively work through this mechanism in DMG.

By immunoblotting, we found that ATRA-induced ADAR degradation in two DIPG/DMG cell lines at least as efficiently as previously reported in pancreatic and breast cancer cell lines (**Fig. 5a**).^38^ Additionally, ATRA markedly reduced proliferation in human DMG cells, whereas untransformed astrocytes were minimally affected, mirroring the effects observed with ADAR KD (**Fig. 5b-d**). Consistent with our findings in human models, dose-dependent ADAR degradation, PD-L1 induction, and gliomasphere killing was also observed in the murine H3K27M cell line NP53,^41, 42^ underscoring the robustness of this response (**Fig. 5e**, **Extended Data Fig. 5a**).

**Figure 5.**
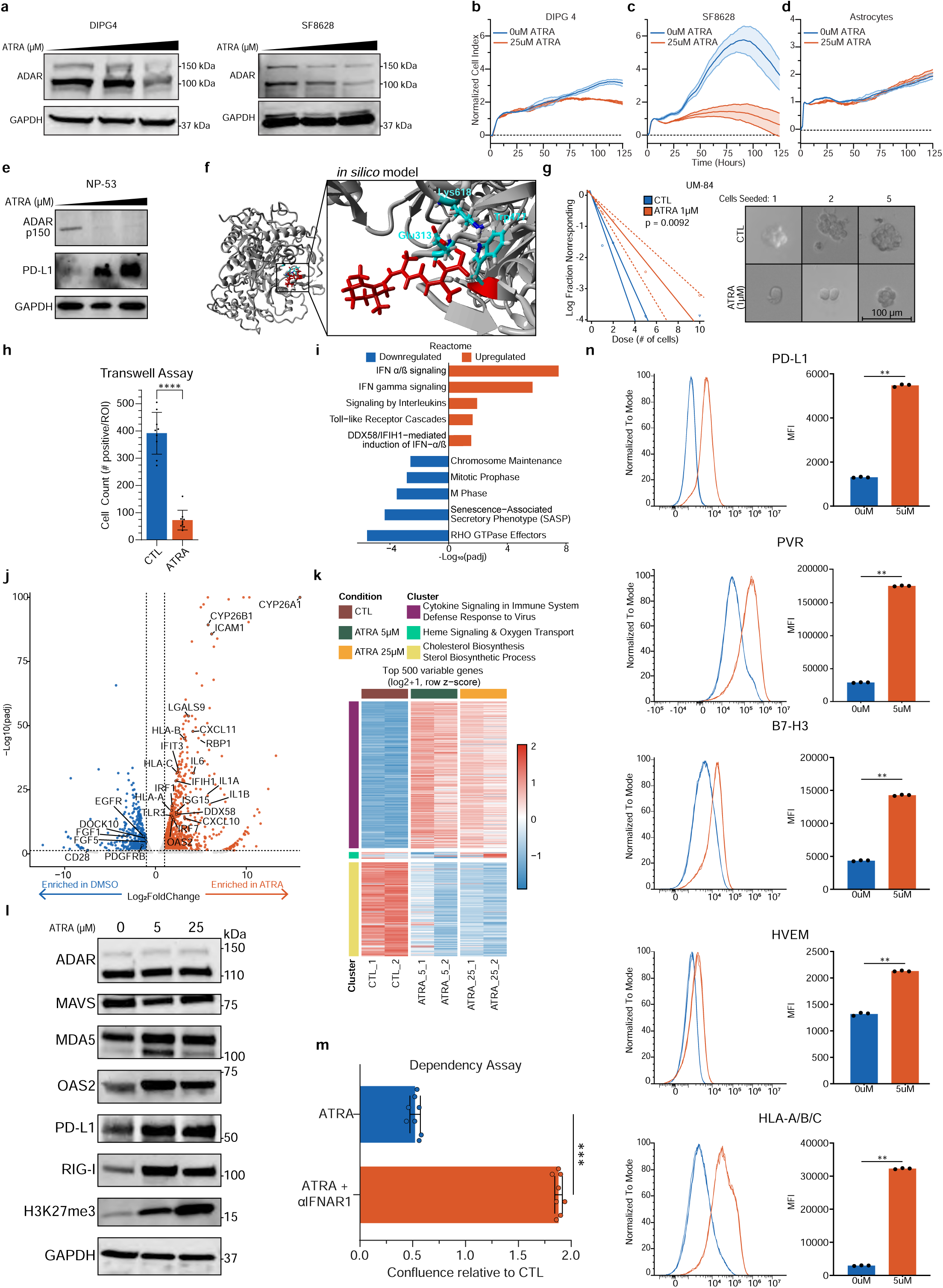
Pharmacologic targeting of ADAR in diffuse midline glioma using all-trans retinoic acid. a) Immunoblot analysis assessing ADAR degradation upon increasing ATRA treatment in two DIPG/DMG cell lines. b-d) xCELLigence real-time proliferation assays demonstrating the effects of 25 µM all-trans retinoic acid on DIPG4 and SF8628 tumor cells compared with untransformed astrocytes. e) Immunoblot assessing ADAR degradation and immune-checkpoint expression upon increasing ATRA treatment in the murine H3K27M cell line NP53. f) *In silico* modeling of ADAR–all-trans retinoic acid interactions elucidates putative binding interfaces. g) Extreme limiting dilution assay evaluating self-renewal capabilities of UM-84 upon 1uM ATRA treatment. h) Assessment of DIPG4 cell migration using a transwell assay following 1 µM ATRA treatment. i) Reactome pathway analysis of 5 µM ATRA-treated cells compared to control. j) Volcano plot comparing significantly differentially expressed genes after 5 µM ATRA treatment compared to control. k) Heatmap of the top 500 differentially expressed genes. Samples are hierarchically clustered by condition, and genes are grouped into three distinct expression clusters based on shared expression patterns, with cluster annotations derived from Enrichr. l) Immunoblot of antiviral response regulators and H3K27me3 marks upon increasing ATRA dosing. m) Dependency rescue assay following 5 µM ATRA using type I interferon receptor blockade in SU-DIPGIV cells. n) Flow cytometric quantification of immune checkpoint ligand expression in response to differing concentrations of ATRA (5, 10, and 25 µM). (H),(M),(N) are presented as mean ± SD;*P < 0.05, **P < 0.01, ***P < 0.001, ****P < 0.0001 and ns: not significant; Chi-squared test for (G), Mann Whitney test for (H),(M), and (N).

Building upon previous work that elucidated ATRA-mediated ADAR degradation is dependent on ubiquitin-proteosome processing,^38^ we performed *in silico* docking analysis to identify the likely conformation by which ATRA binds to ADAR. Using *AutoGridFR*,^43^ the predicted interaction between ADAR and ATRA was dominated by hydrophobic contacts, with a stabilizing hydrogen bond involving Glu313 (**Fig. 5f**). Remarkably, a six-membered ring of ATRA remained solvent exposed, suggesting its functionality as a hydrophobic tag (HyTs) for targeted protein degradation,^44^ or potential for recruitment of a tertiary binding partner, such as an E3 ubiquitin ligase.^45^

Beyond these developments on ADAR-ATRA binding, we sought to understand if *in vitro* treatments with more translationally relevant doses of ATRA (1-5 μM) may activate downstream IFN signaling similar to ADAR KD.

### Translationally relevant dosing of ATRA activates anti-viral signaling and immune activation

Of significant promise, we observed that even at translationally-relevant dosing of ATRA, DMG cells initiated an antitumor response. Similar to siRNA-mediated ADAR KD, human H3K27M cells (UM-84) exhibited significantly decreased self-renewal on ELDA at 1 μM ATRA (**Fig. 5g**). Further, Transwell assays demonstrated significantly reduced migration capacity following 1 µM ATRA treatment (**Fig. 5h**).

To test whether these doses of ATRA may also be activating anti-signaling pathways, we performed RNA sequencing and pathway analysis. Remarkably, pathway analysis (Reactome) revealed significant activation of RIG-I and MDA5 induced IFN signaling and decreased mitosis (**Fig. 5i**). Similarly, KEGG and GO analysis revealed antiviral pathway and other cytokine response upregulation (**Extended Data Fig. 5b, c**). Moreover, in addition to canonical ATRA-induced genes like CYP26A1 and RBP1, several of the genes most differentially expressed aligned with increased cytokine signaling (IL1A, IL1B, CXCL10), antigen presentation (HLA-A, HLA-B, HLA-C), and defense responses to virus (DDX58, IFIH1, OAS2) as well as decreased mitosis and cell proliferation (EFGR, FGF1, PDGFRB) (**Fig. 5j**, **k**). We confirmed upregulation of RLR (MDA5, RIG-I) and OAS2 upon ATRA treatment via immunoblot in the absence of significant ADAR degradation (**Fig. 5l**). Thus, low-dose ATRA treatment appears to upregulate RLR and other dsRNA sensor expression, leading to a robust antiviral response in the dsRNA-overloaded context of DMG, in which these sensors are typically downregulated (**Extended Data Fig. 5d**). Of interest, we also found that H3K27me3 levels positively correlated with ATRA dosing, suggesting that the differentiation effects of ATRA are counteracting the H3K27M-driven epigenetic dysregulation. Notably, and again mirroring ADAR KD, the phenotypic effects of ATRA treatment appeared to be rescued by blocking interferon signaling downstream of RLR activation (αIFNAR1, **Fig. 5m**).

Finally, we asked whether ATRA treatment may synergize with immunotherapy. Following ATRA treatment, we observed significant upregulation of key immune-related cell surface markers, including PD-L1 (**Fig. 5n**). Collectively, our data suggest that, in lieu of a selective ADAR inhibitor, ATRA is a promising adjuvant therapy that activates antiviral signaling against DMG.

### ATRA synergizes with immune-checkpoint inhibition and standard-of-care IR against DMG

Given that ATRA-mediated ADAR degradation enhances tumor immunogenicity and induces immune checkpoint expression, we next sought to determine whether ATRA could function as an effective adjuvant to immunotherapy and standard-of-care irradiation *in vivo*. To evaluate the therapeutic potential of ATRA in DMG, we used the murine H3.3K27M cell line NP53. Cells were cultured under stem-like conditions and used to generate orthotopic genetically engineered mouse models (GEMMs) by stereotactic injection of 200,000 viable cells into the right pons (1 mm lateral, 1 mm ventral, 5 mm depth from the lambda suture). Consistent with prior treatment paradigms for this model,^42^ the treatment period was no longer than 2 weeks, initiated on day 3 after engraftment, with tumor establishment confirmed by IVIS imaging. In addition to overall survival (OS), we monitored tumor growth by IVIS, recorded body weight changes, and performed immunohistochemical analyses on at least two representative tumors per cohort collected at the time of sacrifice.

We first evaluated the efficacy of intraperitoneal ATRA alone (20 mg/kg, three times per week). Although ATRA monotherapy consistently extended survival relative to control, the magnitude of benefit was modest, with a median improvement of approximately one week (**Fig. 6a**). Consistent with these findings, IVIS imaging combined with Ki67 staining demonstrated slower tumor growth in ATRA-treated cohorts (**Extended Data Fig. 6a, b**). No acute toxicities were observed, as assessed by body weight monitoring (**Extended Data Fig. 6c**). Consistent with our *in vitro* findings, immunohistochemical (IHC) analysis of ATRA-treated tumors revealed significantly increased CD8+ T cell infiltration compared with control (**Fig. 6c**, **d**).

**Figure 6.**
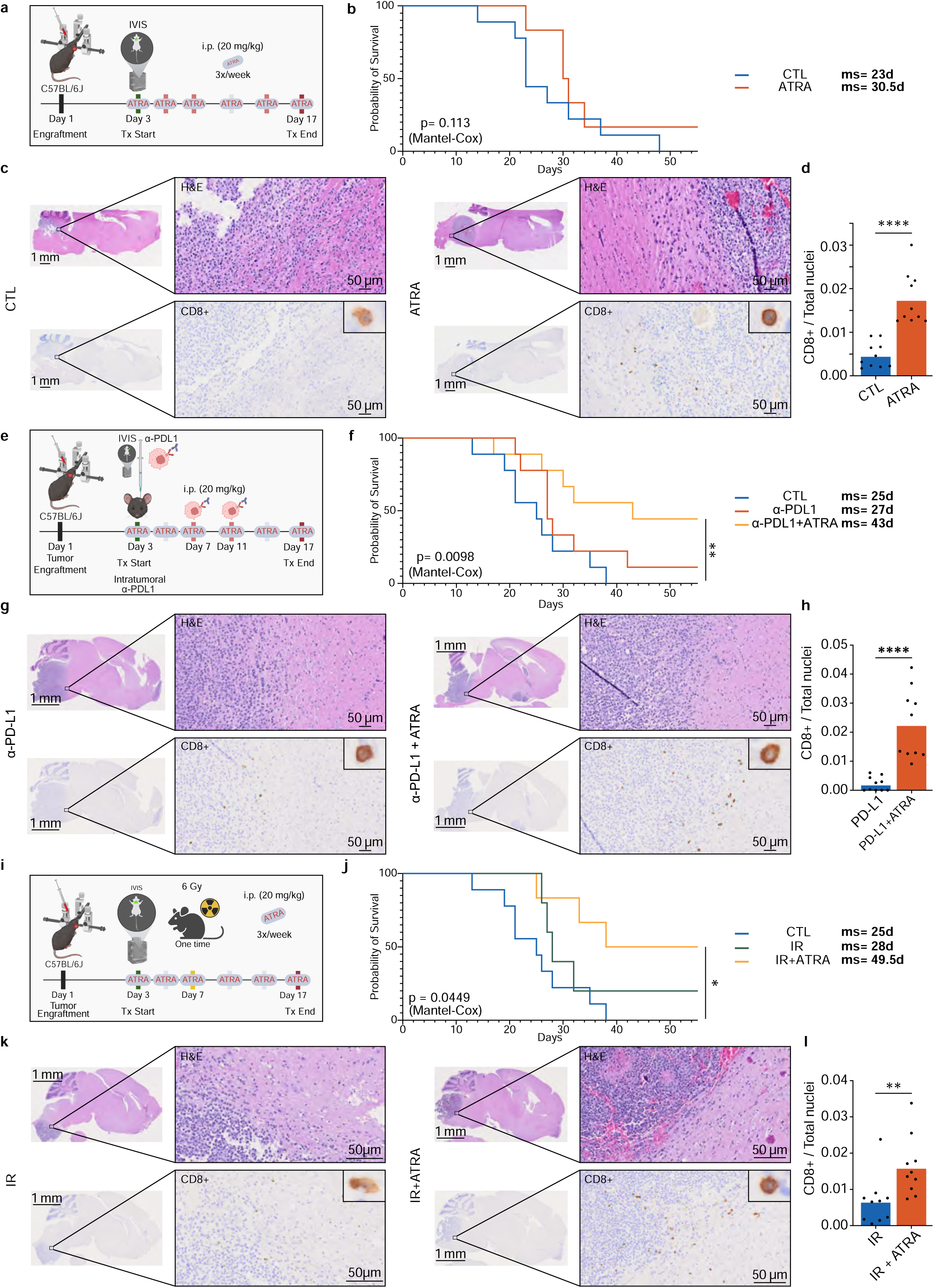
All-trans retinoic acid synergizes with immune checkpoint inhibition and standard-of-care in an in vivo diffuse midline glioma model. a) Schematic illustrating the treatment schedule for the all-trans retinoic acid (ATRA) alone cohort. b) Kaplan–Meier overall survival analysis comparing control (n=9) and ATRA alone (n=6) treatment groups. c, d) Representative hematoxylin and eosin (H&E) staining and CD8 immunohistochemistry (IHC) with quantification of intratumoral CD8⁺ T cell infiltration comparing ATRA-treated tumors with control. e) Schematic illustrating the treatment schedule for the ATRA and intratumoral anti–PD-L1 cohort. f) Kaplan–Meier overall survival analysis comparing control (n=9), anti–PD-L1 alone (n=9), and ATRA+anti-PD-L1 (n=9) treatment groups. g, h) Representative H&E staining and CD8 IHC with quantification comparing tumors treated with ATRA plus anti–PD-L1 versus anti–PD-L1 monotherapy. i) Schematic illustrating the treatment schedule for the ATRA and irradiation (IR) cohort. j) Kaplan–Meier overall survival analysis comparing control (n=9), irradiation alone (n=5), and ATRA+irradiation (n=6) treatment groups. (K), (l) Representative H&E staining and CD8 IHC with quantification comparing tumors treated with ATRA plus irradiation versus irradiation alone. (D),(H),(I), are represented as mean ± SD; *P < 0.05, **P < 0.01, ***P < 0.001, ****P < 0.0001 and ns: not significant; Mann Whitney test for (D), (H) and (I). Mantel-Cox test for (B), (F), and (J).

Despite prior evidence that ATRA penetrates the blood-brain barrier against gliomas^46^ and synergizes with anti-PD1 therapy for pancreatic cancer in vivo and in pilot clinical studies^38^, we observed systemic anti-PD1 therapy (10 mg/kg, twice per week) to be detrimental to the overall survival of DMG mice, in the absence of any acute toxicity (**Extended Data Fig. 6c, d**).

Given the proven safety and encouraging efficacy of locoregional immunotherapy delivery in DMG,^42, 47–49^ we next sought to investigate whether intratumoral (IT) delivery might elicit a more robust therapeutic response. We selected anti–PD-L1 as a more rational agent, as it is abundantly expressed on cancer cells as well as myeloid populations directly accessible within the tumor, whereas PD-1 is primarily only expressed on activated lymphocytes that are relatively sparse in the brain tumor microenvironment.^50^ Using this approach, we observed pronounced synergy between ATRA and intratumoral anti–PD-L1, resulting in significantly slowed tumor growth, a near doubling of OS, and increased CD8+ T cell infiltration (**Fig. 6e-h**, **Extended Data Fig. 6e**). Of note, we did not observe similar effects with regards to systemic anti-PD-L1 therapy (**Extended Data Fig. 6f**).

To enhance the translational relevance of these findings, we lastly assessed whether ATRA synergizes with standard-of-care (SoC) irradiation (IR) (**Figure 6i**). Evaluating ATRA as an adjunct to SoC enables assessment of added therapeutic benefit within a clinically realistic treatment context. This approach allows direct comparison to current practice and supports potential clinical implementation without altering existing treatment paradigms. Accordingly, we observed significant synergistic effects of ATRA and IR in both extending OS by 24.5 days and increasing CD8+ infiltration compared to IR alone (**Fig. 6j-l**, **Extended Data Fig. 6g**).

In summary, ATRA synergizes with both locoregionally delivered anti–PD-L1 therapy and standard-of-care irradiation to significantly improve overall survival and increase CD8⁺ T cell infiltration, supporting its potential utility as an adjuvant therapy in DMG clinical trials.

## Discussion

Despite increased biopsy availability and substantial research efforts, DMGs remain incurable. Previous work has established the H3K27M driver as creating an epigenetically permissive state with repeat dysregulation and therapeutic vulnerability to broad-spectrum epigenetic inhibitors.^13^ In this study, we demonstrate that the H3K27M mutation and its associated epigenetic dysregulation can be leveraged against itself to induce a tumor cell-intrinsic immune response through targeting the RNA editing enzyme ADAR. ADAR KD results in significantly reduced proliferation, migration, and self-renewal. In the absence of FDA-approved agents selectively targeting ADAR or preclinical inhibitors with sufficient brain bioavailability, we established that all-trans retinoic acid (ATRA) is a safe, FDA-approved drug that promotes ADAR degradation at high concentrations. Moreover, antiviral signaling is activated even at translationally relevant ATRA doses, primarily due upregulation of dsRNA sensors, to similarly suppress the proliferative, migratory, and self-renewal properties of DMG cells. Importantly, *in vivo* studies in an immunocompetent model reveal ATRA as a promising adjuvant therapy that can be combined with both immunotherapy and SoC irradiation to extend survival and increase CD8+ infiltration. Taken together, our preclinical data supports the clinical translation of systemically administered ATRA in combination with existing therapeutic modalities and immunotherapy for DMG.

The therapeutic efficacy of immune checkpoint inhibitors has remained limited in aggressive brain tumors such as DMG.^3, 51–53^ Building on our findings that ATRA functions as a potent immunomodulatory agent, we further demonstrate that locoregional delivery of ICIs enhanced the synergistic interaction between ATRA and anti–PD-L1 therapy. While locoregional administration has been a defining feature of recent CAR T cell and oncolytic virus clinical trials in DMG (such as Monje et al, Gállego Pérez-Larraya et al, and Vitanza et al),^48, 49, 54^ ICI clinical trials to date have been administered exclusively IV, including anti-PD-1 (pembrolizumab, NCT02359565; ipilimumab, NCT01952769; and cemiplimab, NCT03690869), anti-PD-L1 (durvalumab, NCT02793466), or anti-CTLA-4+anti-PD-1 (nivolumab + ipilimumab, NCT03130959) regimens.^55^ Based on our data, we propose that future immune checkpoint inhibitor clinical trials for pediatric brain tumors incorporate arms evaluating locoregional delivery, particularly via the intraventricular (ICV) route that may leverage repeated infusions and sampling.

In 2025, the FDA approved ONC-201, the first-ever approved drug for recurrent H3K27M-mutant DMG/DIPG. With this study, we have translational ambitions to evaluate ATRA for rapid integration as an adjunct to the emerging immunotherapy modalities in current Phase 1 DMG clinical trials. We propose that ATRA holds particular promise, as our in vivo data, combined with anti–PD-L1 therapy or irradiation, compare favorably with ONC-201 in H3K27M NP53 models.^56^

In recent years, ATRA has seen a resurgence in clinical trials for solid tumors as an adjuvant to ICI therapy. Of relevance, a Phase II study is currently recruiting recurrent IDH-mutant glioma patients for ATRA and anti-PD-1 therapy (NCT05345002). Future efforts to enhance the translational potential of ATRA will likely depend on overcoming both tumor-intrinsic and microenvironment-mediated protection conferred by CYP26 enzymes, a family of cytochrome P450 monooxygenases that mediate ATRA oxidation and inactivation. Notably, our differential gene expression analysis revealed that *CYP26A1* and *CYP26B1* are among the most strongly upregulated genes following ATRA treatment. This observation aligns with prior work emphasizing the need to combine ATRA with strategies that either protect it or inhibit these retinoic acid–inducible enzymes. Promising approaches include co-delivery of ATRA with small interfering RNA targeting *CYP26A1* via lipid nanoparticles,^57^ as well as concurrent oral administration of ATRA with emerging small-molecule CYP26 inhibitors.^35^ We are continuing to evaluate the combined use of ATRA and CYP26 inhibitors in *in vivo* DMG models to assess both safety and therapeutic efficacy in support of clinical translation.

To our knowledge, the role of epi-transcriptomic regulation in pediatric gliomas has not been previously explored prior to our study. Here, we demonstrate that in the context of H3K27M-driven epigenetic dysregulation, the epi-transcriptomic regulator ADAR functions as a critical checkpoint restraining viral mimicry and innate immune activation. The anti-glioma effects of ADAR inhibition appear to be mediated primarily through the RLR axis, accompanied by concurrent activation of PKR and OAS pathways. In prior work, we demonstrated the preclinical efficacy of RLR–mediated viral mimicry activation in glioblastoma.^58^ Other groups have previously demonstrated ADAR inhibition induces antitumor activity in preclinical glioblastoma models, including reduced tumor growth, remodeling of the tumor microenvironment toward a more immunologically active state, and prolonged survival.^59–61^ However, these studies did not implicate or assess retroelement activation. Building on these findings, we propose epitranscriptomic regulators as key dependencies in epigenetically dysregulated brain tumors, including H3K27M / IDH-mutant gliomas and atypical teratoid/rhabdoid tumors, and highlight neoadjuvant targeting to harness viral mimicry as a strategy to enhance immunotherapy responsiveness in these settings.^23, 62^

## Online Methods

### Experimental model details

#### Mouse models

All animal experiments were approved by the Institutional Animal Care and Use Committee at the University of Miami and performed in accordance with the guidelines. Between 3 and 5 mice were caged together. Standard autoclaved diet and water were provided *ad libitum*. All radiation therapy was performed under isoflurane gas anesthesia, and all efforts were made to minimize suffering.

NP53: 200,000 cells were implanted into 4-week-old C57BL/6J mice (Strain #:000664, The Jackson Laboratory, Bar Harbor, ME, USA) in 3 μl of Dulbecco’s phosphate-buffered saline (DPBS), kindly provided by Dr. Oren Becher (Mount Sinai, NY, USA).^63^ Cells were stereotactically injected into the right pons as previously described in the literature (1 mm right, 1 mm down, 5 mm depth from lambda suture).^42, 64, 65^ After 3 days, mice underwent BLI to confirm tumor uptake and were then randomized to treatment groups. In the event of limited bioluminescence signal three days post-implantation in the optimized diluted model, mice with positive signal were randomized into treatment groups; mice with negative signal were evenly distributed among treatment groups. We ended each experiment and identified long-term survivors to be those animals that lived at least three times longer than the median survival time of control animals.

The NP53 cell line was derived from tumors generated using the RCAS (Replication-competent ASLV long terminal repeat with a splice acceptor) system in NP53 fl/fl mice. These mice were generated by crossing Nestin Tv-a (Ntv-a) mice with p53 floxed mice on a C57BL/6 background (p53 fl/fl). Nestin Tv-a mice express the TVA receptor, which enables RCAS viral infection, under the control of the Nestin promoter. As a result, Nestin-expressing cells are susceptible to viral infection, leading to PDGF-B signaling activation, p53 loss, and ectopic expression of the H3.3 K27 mutation in these cells. Nestin expression is primarily restricted to glial progenitor cells. Dr. Becher derived the NP53 cell line from tumors generated using this system; therefore, these cells are implanted into C57BL/6 mice, the same genetic background in which the tumors were originally generated.

#### Cell lines

SU-DIPG-IV was obtained from Dr. Michelle Monje (Stanford University School of Medicine, Stanford, CA). UM-84 was obtained from Dr. Anna Losorella (University of Miami Miller School of Medicine, Miami, FL). These two cell lines were cultured in serum free neurosphere media containing Neurobasal medium supplemented with GlutaMAX, B27 supplement, N2 supplement, human recombinant EGF, human recombinant bFGF, heparin (2 μg/mL), and penicillin–streptomycin. SF8628 cells was purchased from Millipore Sigma. hTERT-altered astrocytes were a kind gift from Dr. Defne Bayik (University of Miami Miller School of Medicine, Miami, FL). These two cell lines were cultured in DMEM medium DMEM media (Dulbecco’s modified Eagle’s medium (*Thermo Fisher Scientific #11995-065*) supplemented with 10% fetal bovine serum (*Thermo Fisher Scientific #A52567-01*) and 1% antibiotic antimycotic solution (*Thermo Fisher Scientific #A5955-100mL*).

The murine DIPG cell line NP53 (H3.3K27M) was provided by Dr. Oren Becher.^63^ The cell lines were generated from DIPG tumors arising in genetically modified mice. It was cultured in serum free neurosphere media (per 50 mL total volume: 44.5 mL Neurocult basal media (*Stem Cell Technologies*); 10% proliferation supplement (*Stem Cell Technologies #05701);* Pen-strep 1% (*Invitrogen #15140–122*); Human basic FGF, 20ng/mL (*Invitrogen #13256–029)*; Human EGF, 10ng/mL (*Invitrogen #PHG0314)*; Heparin, 2μg/mL (*Stem Cell Technologies #07980)*. Cells were incubated at 37° C and split as required approximately once a week with Accutase (*Stem Cell Technologies*).

#### 293 T-REx Model

Cells were cultured using DMEM media (described above). Stable cell line expressing H3.3K27M transgene were generated using the doxycycline-inducible pINTO-NFH vector expression system. Following transfection, stable integrants were selected using dual-antibiotic selection with zeocin-selectable (200 μg/ml) and blasticidin-selectable (10 μg/ml), stable cell lines were selected through corresponding selectable markers encoded within the pINTO-NFH vector. Sanger sequencing was used to verify successful integration. Doxycycline (1 μg/ml) was added to the culture medium at appropriate time point prior to cell harvest for experimental inductions of the H3.3K27M transgene.

#### RNA Sequencing

Ribosomal depleted RNA was extracted from patient-derived neurosphere cell lysate and sequenced using a lncRNA prep (250–300 bp, paired-end, 50 million reads). RNA-seq was performed by Novogene using the Illumina NovaSeq 6000 PE150 platform. FastQ files were prepared and aligned to the human Hg38 reference genome and subjected to external quality control. A custom bioinformatics pipeline from TE-Transcripts (M. Hammel Laboratory, Cold Spring Harbor Laboratory) was used to annotate retrotransposon counts.^66^

### Bioinformatic analysis

#### Publicly Available Dataset analysis

Publicly available datasets from Clinical Proteomic Tumor Analysis Consortium with the Children’s Hospital of Philadelphia (CPTAC/CHOP), Pediatric Brain Tumor Atlas (PBTA), and The Childhood Cancer Model Atlas (CCMA) were analyzed to characterize transcriptomic and proteomic alterations across low-grade glioma (LGG) and high-grade glioma (HGG) cohorts. When appropriate, datasets were made into subsets to include HGG samples with H3-alterations consistent with DMG diagnosis to ensure molecular uniformity.

#### Clustered Heatmap and Pathway Enrichment Analysis

Gene expression values (FPKM) were log₂-transformed (log₂[FPKM + 1]) and standardized by row-wise z-score scaling. The top 500 most variable genes across samples were selected based on variance and used for downstream analysis. Hierarchical clustering was performed using Euclidean distance and complete linkage, and genes were partitioned into discrete clusters using tree cutting. Genes belonging to each cluster were extracted and exported as individual gene lists. These cluster-specific gene sets were analyzed for functional enrichment using Enrichr. Gene symbols were submitted to the Enrichr web platform, and enrichment results were obtained from the Reactome Pathways 2024 database (Pathways category) and the Gene Ontology (GO) 2025 database (Ontologies category). For each cluster, the most statistically significant enriched terms were selected to represent the dominant biological themes.

#### RNA editing analysis

Adenosine to Inosine (A-to-I) editing events of RNA transcripts were quantified by deploying the REDItools2 pipeline (https://github.com/BioinfoUNIBA/REDItools2) on the institutional load sharing facility. Retroelement editing loci were annotated using hg38 repeat masker accessed via UCSC Genome Browser and analysis of editing counts was performed using R (version 4.4.1) and the Bioconductor (release 3.19) framework. Differential editing count analyses employed DESeq2 (v1.42.1).

#### Survival analysis

Data was accessed from the University of California Santa Cruz, using the PBTA CBTTC dataset. Patients were filtered by histological diagnosis to only include the following: WHO Grade III Glioma (NCIT:C127816), WHO Grade II Glioma (NCIT:C132505), and Diffuse Intrinsic Pontine Glioma (NCIT:C94764). A Kaplan Meier plot was then generated using the site graphical user interface where it split patients by median ADAR1 RSEM TPM gene expression.

#### Western blotting

Total protein was extracted using RIPA lysis buffer (Sigma-Aldrich) with protease and phosphatase inhibitor cocktails (ABCAM, Waltham, MA). Lysates were centrifuged at 14,000 × g for 15 min at 4 °C, and protein concentrations were quantified using the Bio-Rad Protein Assay Reagent according to the manufacturer’s instructions. The protein samples were denatured in a mixture of SDS-PAGE Reducing Agent (×10) and RIPA buffer (×1) at 95 °C for 5 min, then separated on a 4–20% Tris-Glycine SDS-PAGE gels (Bio-Rad), and transferred onto nitrocellulose membranes using a Trans-Blot Turbo transfer system (Bio-Rad). Membranes were blocked for 1 hour at room temperature in 5% Blotting Blotting-Grade Blocker in TBS (Bio-Rad) with 0.1% Tween-20 (TBS-T), then incubated with primary antibodies overnight at 4 °C. Membranes were washed three times with TBS-T followed by incubation with HRP-conjugated anti-rabbit or anti-mouse secondary antibodies (Invitrogen) for 1 h at room temperature. The protein bands were visualized using ECL chemiluminescence (Thermo Fisher Scientific) and imaged with a ChemiDoc imaging system (Bio-Rad). Antibodies utilized in this study can be found in Supplementary Table 2.

#### Silencing RNA (siRNA) transient knockdown

All siRNAs used in this study were obtained from Integrated DNA Technologies (IDT, Coralville, IA). Three siRNAs were initially tested per target, and one candidate was chosen after trialing for degradation by Western blot. Cellular delivery by Lipofectamine RNAiMAX reagent (Thermo Fisher Scientific) according to manufacturer’s instructions. Cells were seeded 24 hours prior to transfection, to achieve ∼60-70% confluency. Cells were collected after 72 hour incubation at 37 °C.

#### Proliferation Assay

Respective cell lines were seeded at 1,000 cells/well in a 96-well E-plate in biological and technical triplicates. Cell proliferation was measured with xCELLigence RTCA DP instrument according to the manufacturer’s instructions (ACEA Bioscience) and visualized for over 7 days in culture or until cell proliferation plateaued.

#### Quantitative PCR (qPCR)

RNA was isolated from cell line samples using the TRIzol extraction protocol. Quantification of RNA was performed using the NanoDrop 2000 (Thermo Fisher Scientific), and the concentration was adjusted to 1000 ng/μL. The RNA was then reverse transcribed using the iScript cDNA Synthesis Kit (1708890). Quantitative PCR (qPCR) was employed to amplify and detect target transcripts on the cDNAs, utilizing 5 μM primers. Detailed information about the specific primers used is provided in Supplementary Table 1.

The qPCR cycling conditions were set as follows: initial denaturation at 95°C for 20 seconds followed by 35 cycles of 95°C for 20 seconds and 60°C for 30 seconds, using the Fast SYBR Green Master Mix. Normalized CT values were used to quantify the expression of target transcripts through the ΔΔCT method. All qPCR runs included reverse transcription negative controls, which did not show amplification. Each qPCR experiment was conducted in both biological and technical triplicates.

#### ELISA

Concentrations of secreted cytokine and chemokine levels were quantified using commercially available IFN-β and CXCL10 ELISA kits (Proteintech) according to manufacturer’s protocol.

Cells were seeded in 6-well plates and either pharmacologically treated or transfected with siRNA as indicated. Culture supernatants were collected 72 hours post-transfection and centrifuged at 1,000 × g for 5 minutes. Clarified supernatants were then analyzed in technical duplicates. Absorbance was measured at 450 nm wavelength using a microplate reader. Data was analyzed using GraphPad Prism (v.11; GraphPad Software).

#### In vitro all-trans retinoic acid (ATRA) treatment

All-trans retinoic acid (ATRA; Millipore Sigma, PHR1187-3X100MG) was dissolved in dimethyl sulfoxide (DMSO) to generate a 10 mM stock solution and stored according to the manufacturer’s recommendations. For in vitro experiments, the stock solution was diluted in complete culture medium to a final concentration of 5 µM immediately prior to use. Cells were treated with 5 µM ATRA for 72 hours, with fresh ATRA-containing medium replaced every 24 hours to maintain compound stability and activity. Control cells receive an equivalent volume of DMSO. After 72 hours of treatment, cells were harvested for downstream analyses.

#### Flow cytometry

Cell-surface immunophenotyping was performed on patient-derived DMG cell SU-DIPG-IV. Cells were collected 72 hours after siRNA transfection and washed twice with phosphate-buffered saline (PBS) and resuspended in fluorescence-activated cell sorting (FACS) buffer (PBS supplemented with 2% fetal bovine serum and 2 mM EDTA). To exclude dead cells, samples were stained with a fixable viability dye and incubated for 15 minutes, following manufacturer’s instructions. Cells were then stained with antibody panels, as listed in Supplementary Table 2.

#### RNA immunoprecipitation (RIP)

After cell harvesting, cytoplasmic fractions were extracted using the Nuclear Extract Kit (Active Motif; no. 40010) according to the manufacturer’s instructions. To isolate RNA, an equal volume of 70% ethanol was added to the cytoplasmic fractions, and RNA purification was performed using the TRIzol method followed by DNAse digestion. The total RNA was dissolved in 38 μL RNase-free water. A total 2 microliters of total RNA were used as input, and the remaining RNA was divided into 2 tubes.

#### Dependency Assay

SU-DIPG-IV (DIPG4) cells were seeded at 0.5×10^5 cells per well in 6-well plates.

Control or ADAR-targeting siRNA duplexes were diluted in Opti-MEM reduced-serum medium (Thermo Fisher Scientific), per manufactures instructions. PKR was inhibited using a small molecule PKR inhibitor (Sigma Aldrich) (250 nM), while Type I interferon signaling was blocked using an anti-IFNAR1 ((BioXcell) 10 µg/mL). Experimental conditions tested included control siRNA, ADAR knockdown (KD), ADAR KD + PKRi, and ADAR KD + anti-IFNAR1. After 72 hours, cells were then collected and analyzed for culture density.

#### Crystal Violet Staining Assay

Cells were cultured in biological duplicate under the following conditions: siCTL, siADAR, siADAR + ∝IFNAR1, and siADAR+PKRi for 72 hours prior to analysis. Culture density was subsequently assessed using a Crystal Violet Assay Kit (Abcam ab232855) in accordance with the manufacturer’s protocol. After removal of culture medium, cells were washed with 1x Wash Buffer and stained with crystal violet solution containing methanol for 20 minutes at room temperature. Following incubation, excess dye was removed, and plates were thoroughly washed with 1x Wash Buffer I. Plates were then air-dried prior to imaging. Cell growth was quantified using at least 4 representative Brightfield images per well and percentage confluency was determined using EVOS analysis software (Thermo Fisher Scientific). Data and statistical analysis were performed using GraphPad Prism.

#### Transwell Migration Assay

Cell migration was assessed using Transwell 3.0 µm Polyester membrane inserts (Corning). Medium supplemented with 2.5% fetal bovine serum (FBS) served as a chemoattractant solution in the lower chamber. Transwell inserts were removed from the plate and placed into the wells containing the chemoattractant, and serum-free medium containing DMG cells were added to the upper chamber. To allow for migration, cells were incubated for 24 hours at 37 °C in humidified 5% CO₂. Following incubation, 5% paraformaldehyde (PFA) was added to the lower wells for 5 minutes to fix migrated cells on the underside of the inserts in PFA. Non-migrated cells on the upper surface were gently removed using a cotton swab, with the lower surface left intact. Inserts underwent two sequential 5-minutes wash in phosphate-buffered saline (PBS). Migrated cell nuclei were stained with DAPI or Hoechst (1:1000 dilution) and incubated for 5 min. After which, membranes were excised using a scalpel, mounted onto glass slides with mounting medium, and sealed with coverslips. The underside of the migrated cells membrane were visualized using an EVOS fluorescence microscope (Invitrogen). Quantification was performed by counting stained nuclei in representative fields of view using EVOS analysis software. Migrated cells were quantified for each condition from three independent replicates. Statistical analyses between groups were performed in GraphPad Prism (v.11; GraphPad Software), applying an unpaired, two-tailed Student’s T-test, with p < 0.05 considered statistically significant.

#### Extreme Limiting Dilution Assay (ELDA)

DMG neurospheres were enzymatically dissociated into single cell suspension by incubation in TrypLE solution for 5 minutes at 37 °C. Cells were counted using a hemocytometer and plated into a 96-well ultra-low attachment plate at various serial dilutions (1,2,5, or 10 cells per well). Twenty-four replicate wells were prepared for each seeding density, and the dilution series was repeated across the plate to maximize statistical power. All wells were cultured in neurosphere medium and maintained in incubation at 37 °C with 5% CO₂ for 7 to 14 days. Every three days after plating, wells were examined by phase-contrast microscopy using an EVOS imaging system, and images were acquired. Over the course of day 7 through day 14, wells were classified as positive if they contained at least one neurosphere with a diameter of *20 µm* or greater. Sphere diameters were quantified using the calibrated measurement tool within the imaging software. For each seeding density, the number of positive wells were recorded and analyzed using the ELDA software package (http://bioinf.wehi.edu.au/software/elda) with the default 95% confidence interval. Graphical outputs were exported in PDF format, and numerical results were compiled for quantitative analysis and visualization.

#### Immunohistochemistry (IHC)

IHC was performed using the BOND Refine Polymer Detection Kit (Leica Biosystems, cat #NC0318637) on the Leica BOND RXm automated research stainer (Leica Biosystems, Germany) per manufacture’s protocol. Antigen retrieval was performed with BOND ER Solution 1 (citrate buffer, pH 6.0) for 20 minutes at 100°C followed by four Bond Wash Solution washes. Peroxide Block was used to block endogenous peroxidase activity for 5 minutes and washed three times with Bond Wash solution. To prevent non-specific binding, Slides were blocked with 10% normal goat serum for 30 minutes at room temperature was used. Slides were then incubated for 30 minutes with primary antibodies, CD8 (Cell Signaling Technology, cat. #98941S) and Ki67 (Cell Signaling Technology, cat. #12202S) followed by 6 Bond Wash Solution washes. Signal was developed using mixed 3,3’-Diaminobenzidine (DAB) refine reagent for 10 minutes, after which slides were washed with distilled water three times. Slides were then stained with hematoxylin for 5 minutes and rinsed again with distilled water three times followed by three Bond Wash Solution washes and given one final distilled water rinse prior to cover slipping.

## Extended Data Figure Legends

**Extended Data Figure 1, ADAR upregulation in DMG.**

a. Interferon stimulated gene (ISG) scores between H3K27M HGG tumor biospecimens compared to WT HGGs and other pediatric brain tumors (data from the Pediatric Brain Tumor Atlas, n=455).
b. ADAR copy number gain percentage in H3K27M mutant pHGGs to WT (n=20).
c. ADAR RNA levels across pediatric cell lines (data from the Cancer Cell Model Atlas, n=54).
d. represented as a violin plot where the central dashed line is the median and the upper and lower quartiles are the dotted lines above and below, respectively. (C) represented as a box and whisker plot where the central line is the median and upper and lower quartiles are the edges of the box; Ordinary one-way ANOVA for (A) and (C) and with pairwise T-test for (A); *P < 0.05, **P < 0.01, ***P < 0.001, ****P < 0.0001 and ns: not significant;

**Extended Data Figure 2, Effects of ADAR KD in DMG.**

a. siRNA assessment in DIPG4 from three candidate sequences.
b. Representative images following ADAR KD for extreme limiting dilution assay (ELDA) in UM-84
c. Representative images following ADAR KD for transwell migration assay in DIPG4.
d. Representative images following ADAR KD and subsequent rescue with either IFNAR1 blockade or PKR inhibitor (dependency assay) in DIPG4.
e. Immunoblot of ADAR (CRISPR) knockout compared to non-targeting sgRNA DIPG4 cell lines shows adaptive expression of several key viral mimicry regulators and the immune checkpoint ligand PD-L1.

**Extended Data Figure 3, Viral mimicry activation upon ADAR KD.**

a. Reactome pathway analysis (Upregulated, right; downregulated, left) for ADAR KD compared to siCTL in DIPG4.
b. qPCR result for retrotransposon levels by class and representative interferon-stimulated gene under ADAR KD and siCTL conditions in DIPG4.
c. (B) represented as mean ± SD;*P < 0.05, **P < 0.01, ***P < 0.001, ****P < 0.0001 and ns: not significant; Unpaired two-tailed t-test for (B).

**Extended Data Figure 4, H3K27M+ synthetic immunogenicity upon ADAR KD**

a. qPCR result for representative interferon-stimulated gene under ADAR KD and siCTL conditions in both doxycycline-induced and uninduced TREX cells.
b. qPCR result for retrotransposon levels by class under ADAR KD and siCTL conditions in both doxycycline-induced and uninduced TREX cells and (B) represented as mean ± SD;*P < 0.05, **P < 0.01, ***P < 0.001, ****P < 0.0001 and ns: not significant; Kruskal-Wallis test for (A) and (B).

**Extended Data Figure 5, ATRA-induced antiviral signaling in DMG.**

a. Representative images and quantification (stratified by gliomasphere diameter as proxy for size) for gliomasphere formation assay under increasing ATRA dosing in NP53 murine cell line.
b. KEGG pathway analysis for 5 µM ATRA compared to control in DIPG4.
c. GO pathway analysis for 5 µM ATRA compared to control in DIPG4. (A) represented as stacked boxplot with mean ± SD.

**Extended Data Figure 6, ATRA synergy with immune checkpoint inhibitors and irradiation.**

a. Bioluminescence monitoring illustrates tumor kinetic differences between control, ATRA, and α-PD1, and combination treatment.
b. Representative hematoxylin and eosin (H&E) staining and immunohistochemistry (IHC) for Ki67 between control, ATRA, and α-PD1, and combination treatment cohorts.
c. Weight changes at timed intervals after control, ATRA, α-PD1, and combination treatment.

anti–PD-1 alone (n=5), and combination (n=8)

1. d) Schematic and Kaplan–Meier overall survival analysis comparing control (n=9), ATRA alone (n=6), anti–PD-1 alone (n=5), and combination (n=8).
2. e) Bioluminescence monitoring illustrates tumor kinetic differences between control, intratumoral α-PDL1, and α-PDL1+ATRA treatment.
3. f) Kaplan–Meier overall survival analysis comparing control, systemic anti–PD-L1 alone, and combination treatment groups.
4. g) Bioluminescence monitoring illustrates tumor kinetic differences between control, ATRA, irradiation, and combination treatment. (A),(C) represented as mean ± SD; *P < 0.05, **P < 0.01, ***P < 0.001, ****P < 0.0001 and ns: not significant; Mantel-Cox test for (D).

## Acknowledgments

We thank the shared resources facilities at the University of Miami Miller School of Medicine (Cancer Modeling Shared Resource) for support.

## Author contributions

C.K.R. conceived the study. D.R. and A.H.S. provided overarching supervision and strategic direction. C.K.R. led the experimental design under the guidance of the supervising authors. Methodology and experimental procedures were carried out by C.K.R., V.A., F.S., K.K., M.A., and D.S. The principal investigative work was conducted by C.K.R. with additional contributions from the authors listed above. C.K.R. and F.S. performed all bioinformatic analyses and generated the visualizations and publication-quality figures. Data curation and formal analysis were undertaken by C.K.R., F.S., and D.S. C.K.R. drafted the manuscript, and all authors contributed to critical revision and approved the final version.

## Data availability

Gene expression datasets will be deposited to GEO upon publication.

## Code availability

Not applicable.

