## Extended Data Figures 1-6 for "Targeting an RNA Editor to Impede H3K27M+ Pediatric Gliomas"

### Extended Data Figure 1

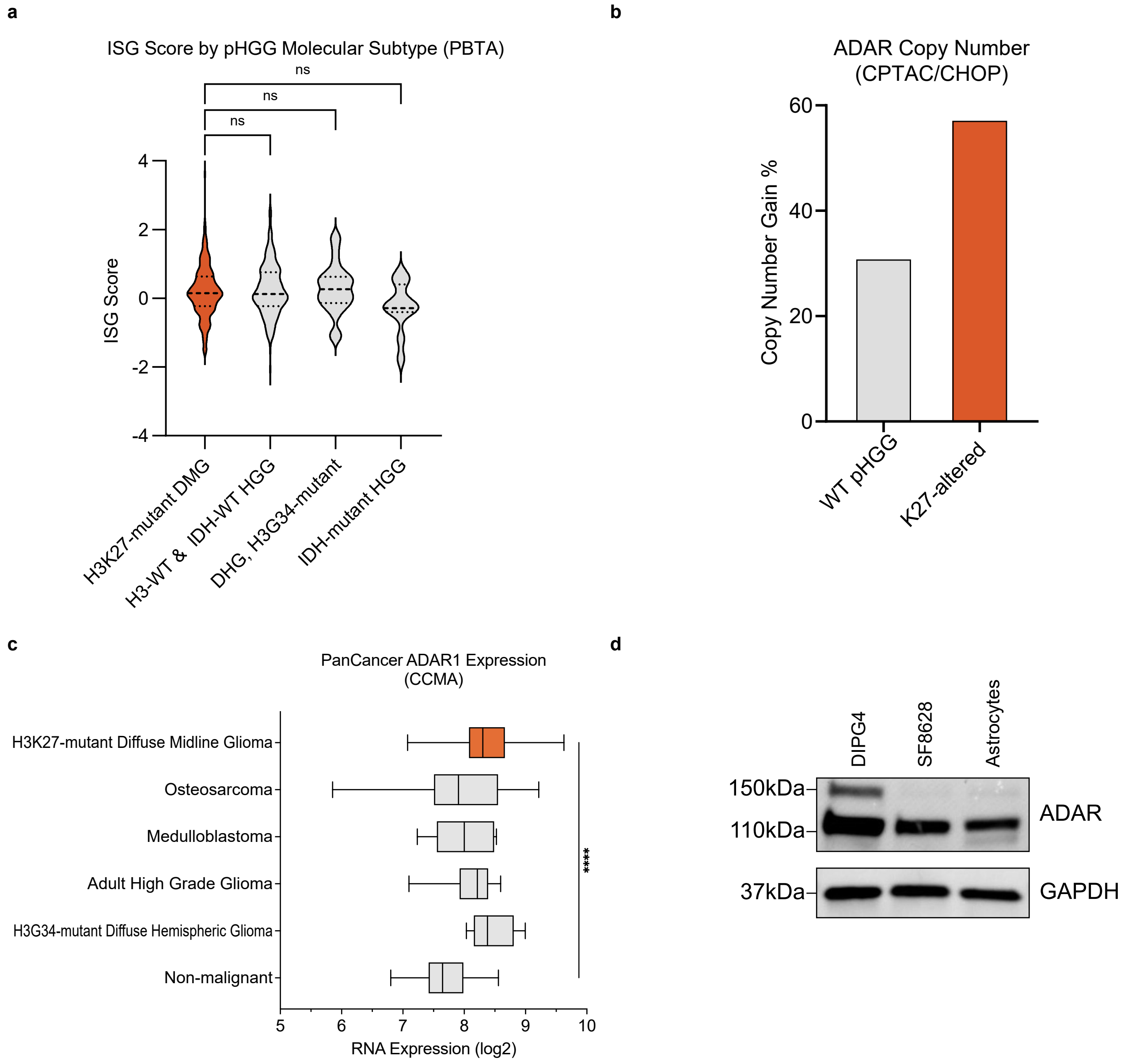

Extended Data Figure 2

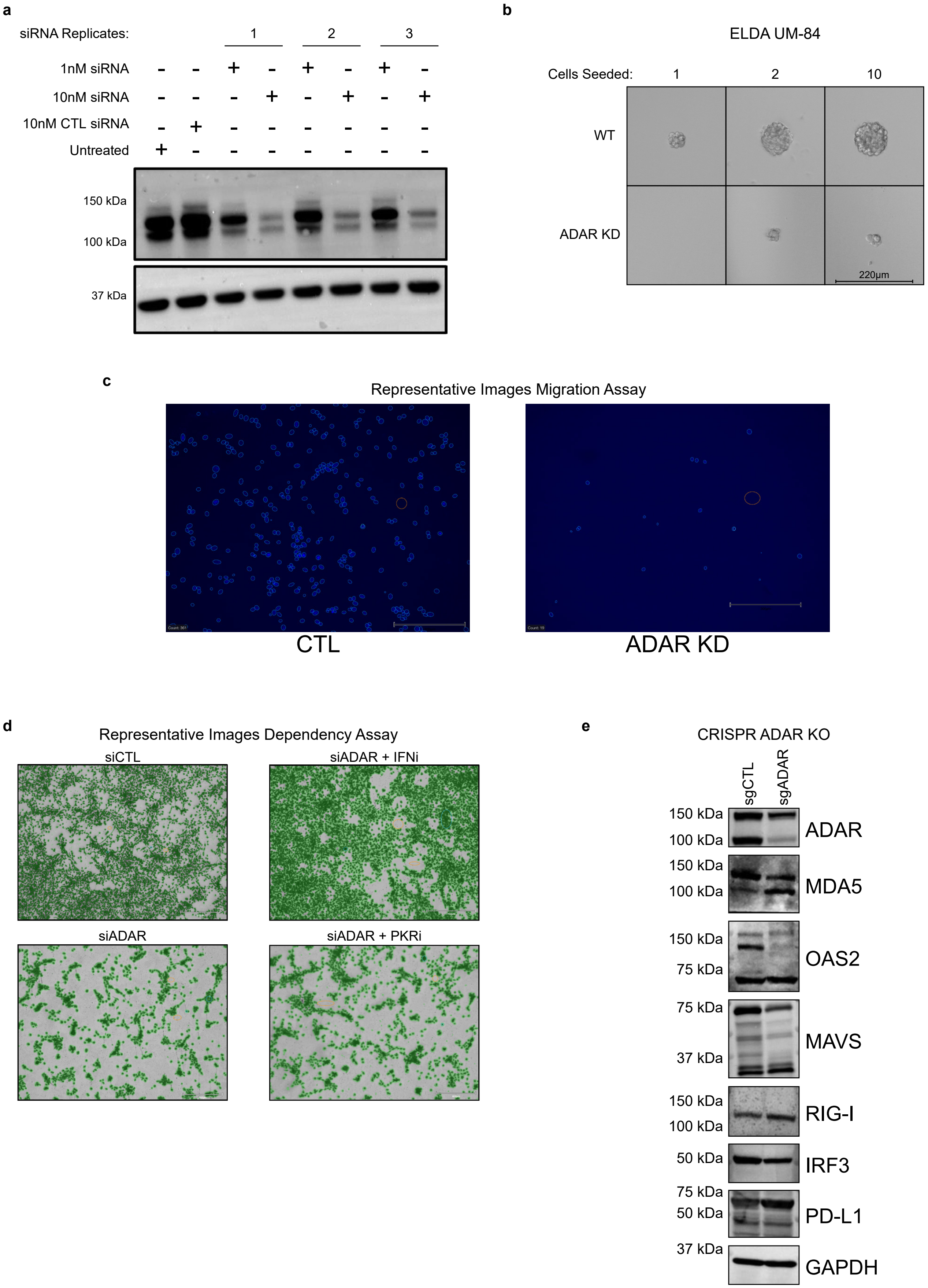

### Extended Data Figure 3

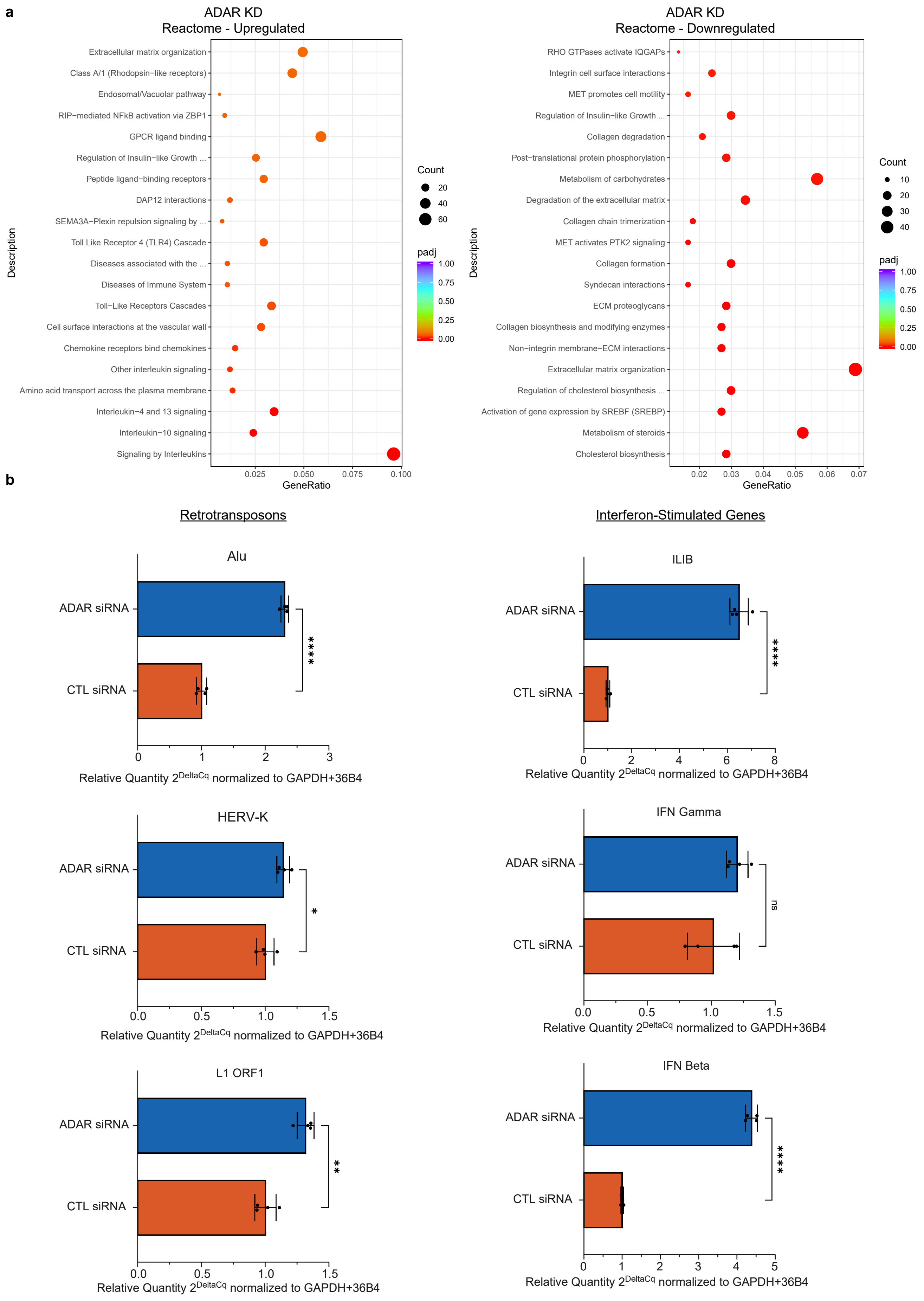

Extended Data Figure 4

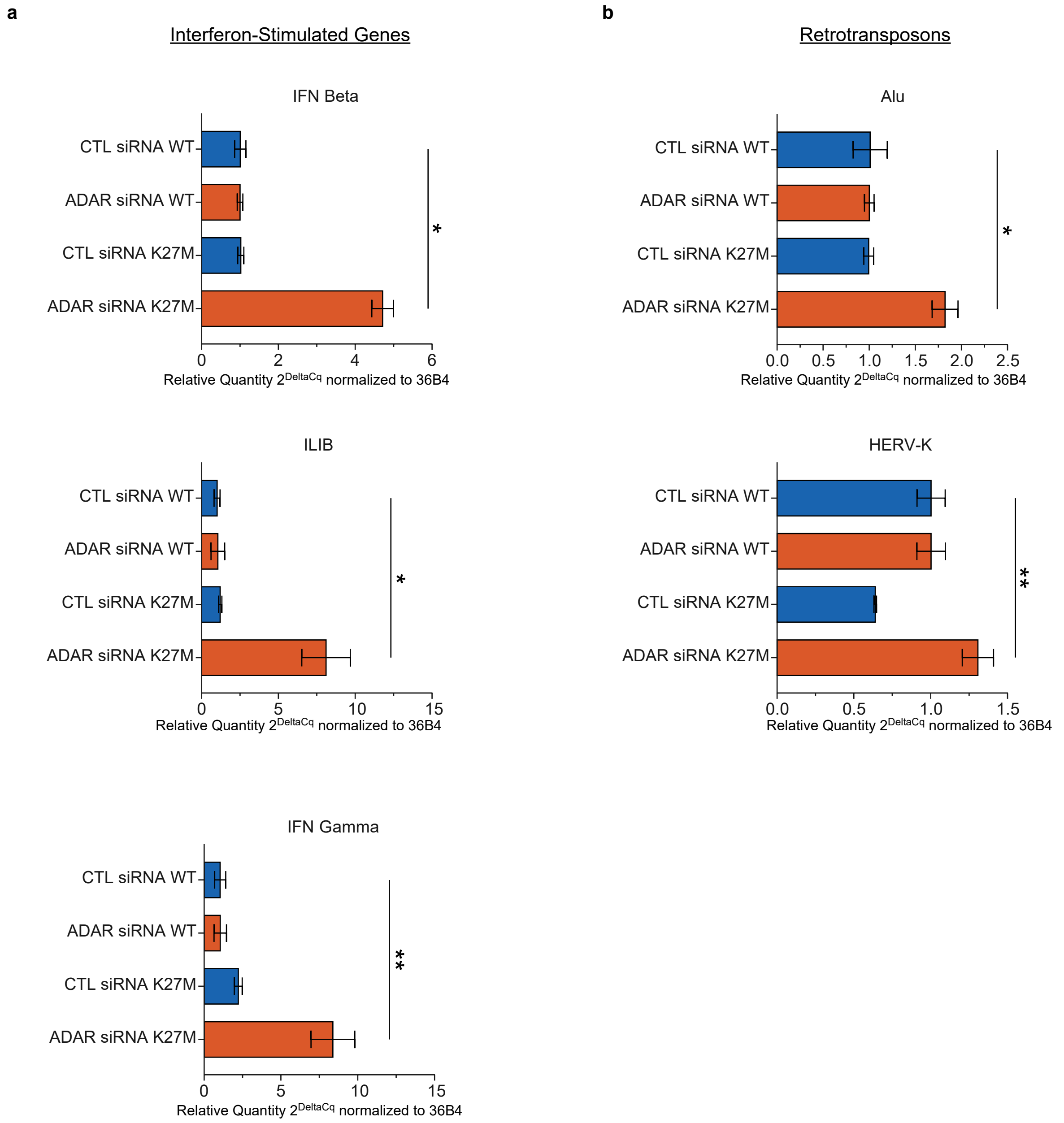

### Extended Data Figure 5

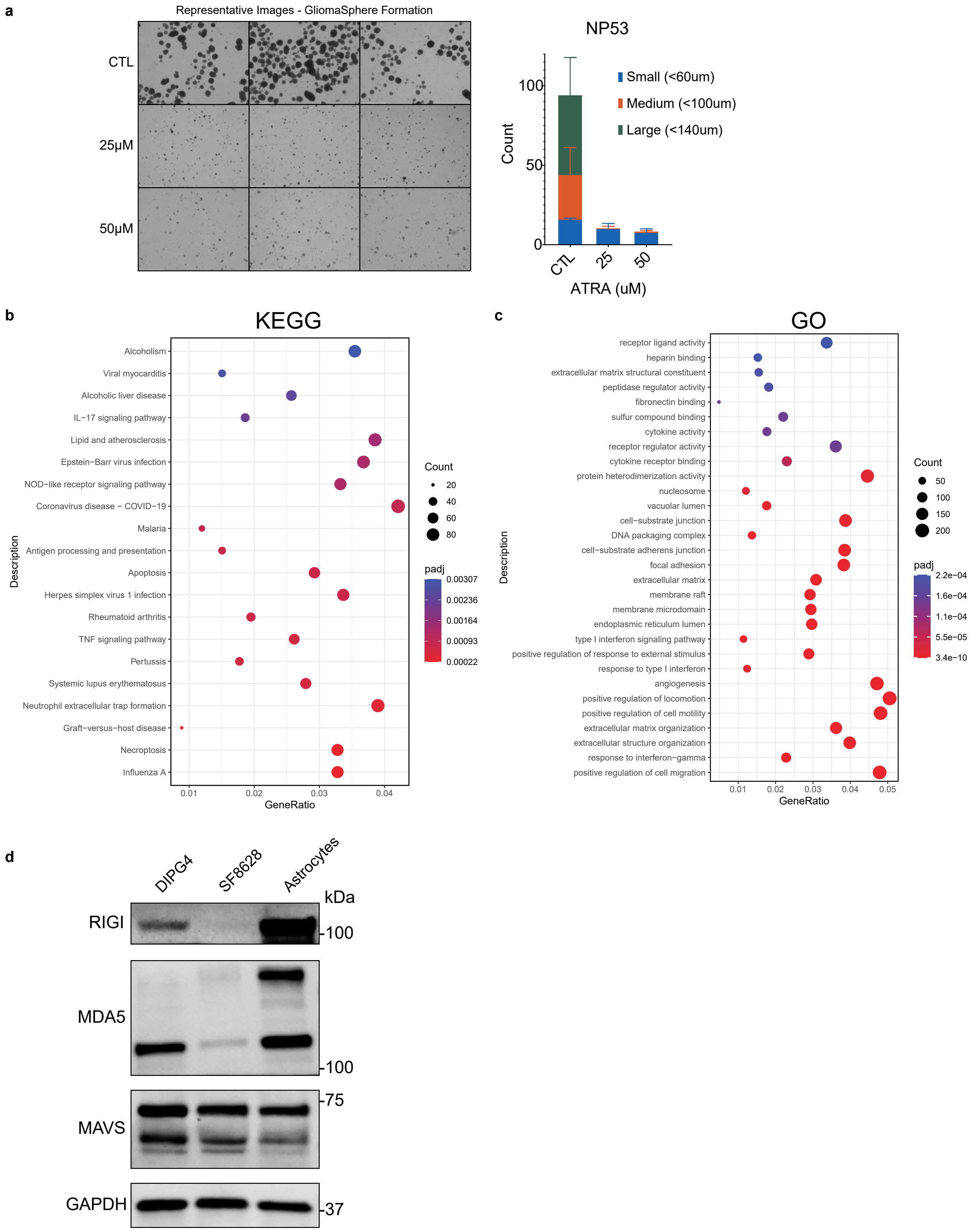

Extended Data Figure 6

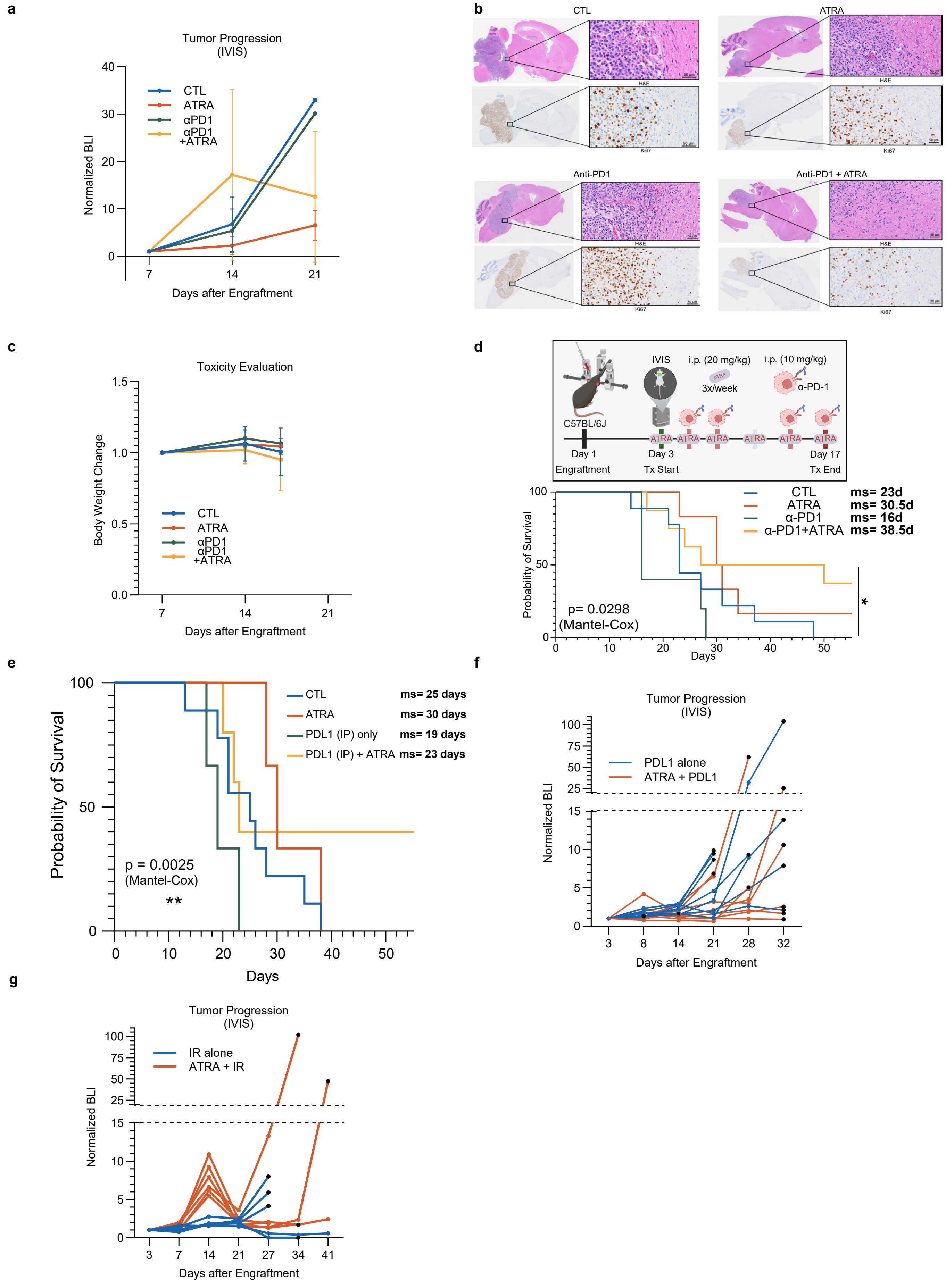
